# Loss of CTRP10 results in female obesity with preserved metabolic health

**DOI:** 10.1101/2023.11.01.565163

**Authors:** Fangluo Chen, Dylan C. Sarver, Muzna Saqib, Leandro M Velez, Susan Aja, Marcus M. Seldin, G. William Wong

## Abstract

Obesity is a major risk factor for type 2 diabetes, dyslipidemia, cardiovascular disease, and hypertension. Intriguingly, there is a subset of metabolically healthy obese (MHO) individuals who are seemingly able to maintain a healthy metabolic profile free of metabolic syndrome. The molecular underpinnings of MHO, however, are not well understood. Here, we report that CTRP10/C1QL2-deficient mice represent a unique female model of MHO. CTRP10 modulates weight gain in a striking and sexually dimorphic manner. Female, but not male, mice lacking CTRP10 develop obesity with age on a low-fat diet while maintaining an otherwise healthy metabolic profile. When fed an obesogenic diet, female *Ctrp10* knockout (KO) mice show rapid weight gain. Despite pronounced obesity, *Ctrp10* KO female mice do not develop steatosis, dyslipidemia, glucose intolerance, insulin resistance, oxidative stress, or low-grade inflammation. Obesity is largely uncoupled from metabolic dysregulation in female KO mice. Multi-tissue transcriptomic analyses highlighted gene expression changes and pathways associated with insulin-sensitive obesity. Transcriptional correlation of the differentially expressed gene (DEG) orthologous in humans also shows sex differences in gene connectivity within and across metabolic tissues, underscoring the conserved sex-dependent function of CTRP10. Collectively, our findings suggest that CTRP10 negatively regulates body weight in females, and that loss of CTRP10 results in benign obesity with largely preserved insulin sensitivity and metabolic health. This female MHO mouse model is valuable for understanding sex-biased mechanisms that uncouple obesity from metabolic dysfunction.

## INTRODUCTION

The prevalence of obesity has nearly tripled in the past four decades and the underlying cause is complex and multifactorial (1, 2). Genetics, environmental and social factors, and demographics all play a role in contributing to excessive weight gain in the setting of overnutrition (3, 4). Although obesity is a major risk factor for type 2 diabetes, dyslipidemia, cardiovascular disease, and hypertension, not all obese individuals develop the metabolic syndrome (5). There is a subset of metabolically healthy obese (MHO) individuals with an apparently healthy metabolic profile free of some or most components of the metabolic syndrome (6, 7). The molecular and physiological underpinnings of MHO are, however, not well understood. Novel preclinical animal models that can recapitulate features of MHO will be valuable in illuminating pathways that resist the deleterious effects of obesity and provide new therapeutic avenues to mitigate obesity-linked comorbidities.

The mechanisms that normally maintain body weight and metabolic homeostasis are complex and involve both cell autonomous and non-cell autonomous mechanisms. Tissue crosstalk mediated by paracrine and endocrine factors plays an especially important role in coordinating metabolic processes across organ systems to maintain energy balance (8). Of the secretory proteins that circulate in plasma, C1q/TNF-related proteins (CTRP1-15) have emerged as important regulators of insulin sensitivity, and glucose and lipid metabolism (9). We originally identified the first seven members of the CTRP family based on shared sequence homology to the insulin-sensitizing adipokine, adiponectin (10), and subsequently characterized eight additional members (11–16). All fifteen CTRPs share a common C-terminal globular C1q domain and are part of the much larger C1q family (17, 18). The use of gain- and loss-of-function mouse models has helped establish CTRP’s role in controlling various aspects of sugar and fat metabolism (12, 19–34). Additional diverse functions of CTRPs have also been demonstrated in the cardiovascular (35–47), renal (48, 49), immune (20, 50, 51), sensory (52, 53), gastrointestinal (54), musculoskeletal (55–57), and the nervous system (58–61).

Of the family members, CTRP10 (also known as C1QL2) is understudied and consequently only limited information is available concerning its function. The best characterized role of CTRP10 is in the central nervous system (CNS). It has been shown that CTRP10 secreted from mossy fibers is required for the proper clustering of kainite-type glutamate receptors on postsynaptic CA3 pyramidal neurons in the hippocampus (62). It serves as a transsynaptic organizer by directly binding to Neurexin3 (Nrxn3) on the presynaptic terminals, and to glutamate ionotropic receptor kainate type subunit 2 (GluK2) and GluK4 on the postsynaptic terminals (62). Additional putative roles of CTRP10 in the CNS have also been suggested. Genome-wide association studies (GWAS) have implicated CTRP10/C1QL2 in cocaine use disorder (63). In rat models of depression, *Ctrp10* expression is increased in the dentate gyrus and reduced in the nucleus accumbens (64). In humans with a history of psychiatric disorders (e.g., schizophrenia), the expression of *CTRP10* is elevated in the dorsolateral prefrontal cortex of both males and females (64). Whether and how CTRP10 contributes to addictive behavior and psychiatric disorders is unknown. Recent large-scale efforts to illuminate the genetic architecture of the human plasma proteome also highlighted an association of plasma C1QL2/CTRP10 levels with a missense C4BPA variant (65). In pregnancy, circulating C1QL2/CTRP10 levels are negatively associated with the plasma inflammasome marker, α1-acid glycoprotein (AGP) (66). The biological significance of these clinical observations, however, remains unclear.

The potential function of CTRP10 in peripheral tissues, however, is essentially unknown and unexplored. The present study was motivated by the well documented metabolic functions of many CTRP family members we have characterized to date using genetic loss-of-function mouse models (19–28, 30, 31). We determined that the expression of *Ctrp10* in peripheral tissues is modulated by diet and nutritional states, and thus may have a metabolic role. We therefore used a genetic loss-of-function mouse model to determine if CTRP10 is required for regulating systemic metabolism. We unexpectedly discovered a female-specific requirement of CTRP10 for body weight control. We showed that the *Ctrp10* KO mice represent a unique female model of MHO with largely preserved insulin sensitivity and metabolic health. This valuable mouse model can be used to inform sex-dependent mechanisms that uncouple obesity from insulin resistance, dyslipidemia, and metabolic dysfunction.

## RESULTS

### Nutritional regulation of *Ctrp10* expression in the brain and peripheral tissues

CTRP10 protein is highly conserved from zebrafish to human (Fig. 1A), with amino acid identity of 67%, 71%, 77%, and 94% between the full-length human protein and the fish, frog, chicken, and mouse orthologs, respectively. The conservation is much higher at the C-terminal globular C1q domain (93-100% identity) between the orthologs. Among the 12 different mouse tissue examined, brain had the highest expression of *Ctrp10* (Fig. 1B), consistent with previous findings (67). Expression of *Ctrp10* in peripheral tissues was variable and generally much lower than the brain (Fig. 1B). We first determined whether *Ctrp10* expression is modulated by nutrition and metabolic state. Male mice (∼10 weeks old) were subjected to fasting and refeeding. In the refed period after an overnight fast, we observed a significant downregulation of *Ctrp10* in the visceral (gonadal) white adipose tissue (gWAT), liver, skeletal muscle, kidney, cerebellum, cortex, and hypothalamus relative to the fasted state (Fig. 1C). Next, we examined whether an obesogenic diet alters the expression of *Ctrp10*. Male mice fed a high-fat diet (HFD) for 12 weeks had a modest increase in *Ctrp10* expression in brown adipose tissue (BAT) and heart and decreased expression in skeletal muscle relative to mice fed a control low-fat diet (LFD) (Fig. 1D). These data indicate that *Ctrp10* expression is dynamically regulated by acute alterations in energy balance, and perhaps to a lesser extent in alterations to chronic nutritional state on an obesogenic diet.

**Figure 1.**
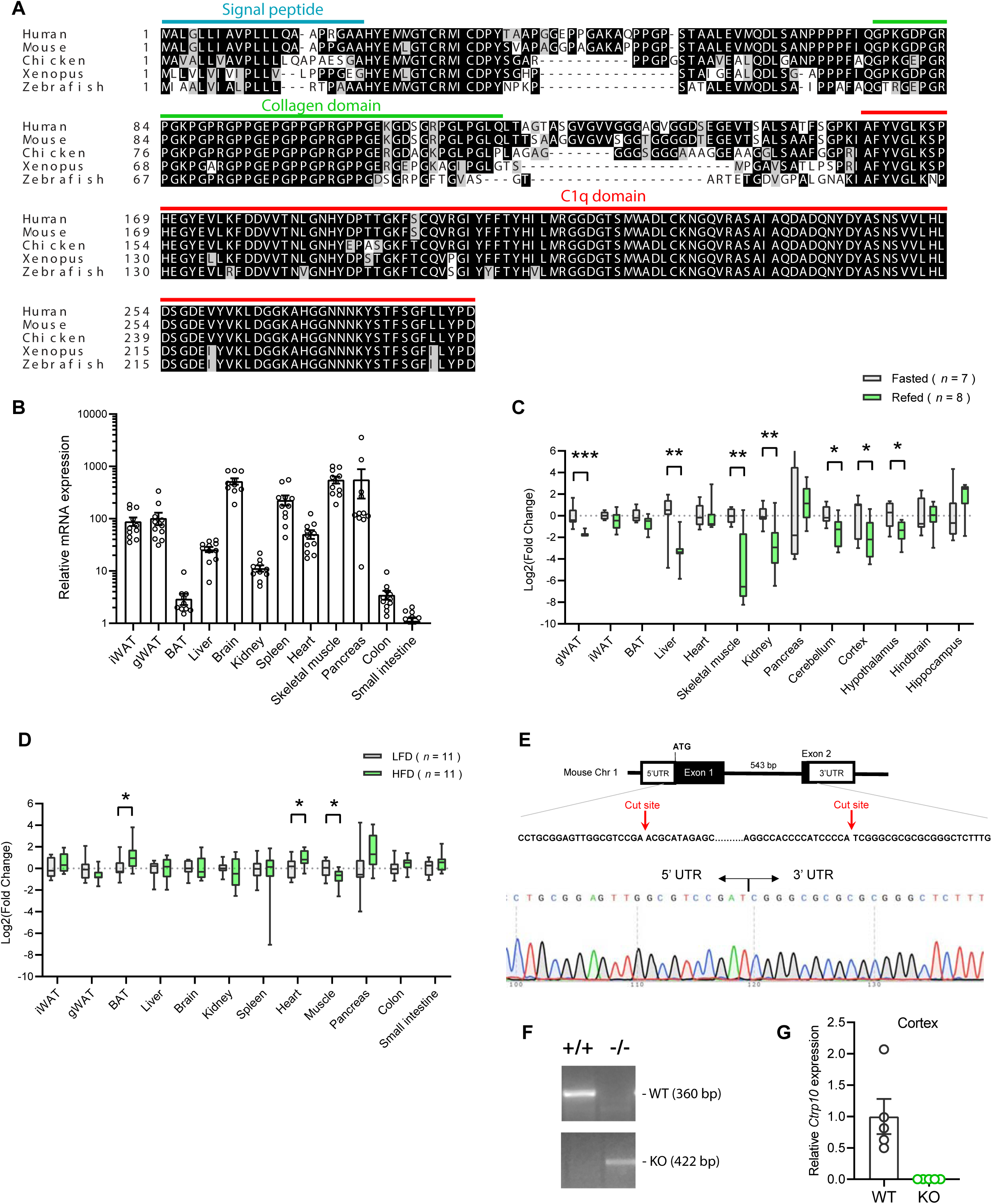
Nutritional regulation of *Ctrp10* expression. **(A)** Sequence alignment of full-length human (GenBank # NP_872334), mouse (NP_997116), chicken (XP_046777733), xenopus frog (XP_031749381), and zebrafish (XP_001920705) CTRP10/C1ql2 using Clustal-omega (141). Identical amino acids are shaded black and similar amino acids are shaded grey. Gaps are indicated by dash lines. Signal peptide, collagen domain with characteristic Gly-X-Y repeats, and the C-terminal globular C1q domain are indicated. **(B)** *Ctrp10* expression across different mouse tissues (*n* = 10). **(C)** Expression of *Ctrp10* across mouse tissues in response to an overnight (16 h) fast or fasting followed by 2 h refeeding. **(D)** Expression of *Ctrp10* across mouse tissues in response to a high-fat diet (HFD) for 12 weeks or a control low-fat diet (LFD). **(E)** Generation of *Ctrp10* knockout (KO) mice. The entire protein coding region in exon 1 and 2 of *Ctrp10* was deleted using CRISPR/Cas9 method and confirmed with DNA sequencing. **(F)** Wild-type (WT) and KO alleles were confirmed by PCR genotyping. **(G)** The complete loss of *Ctrp10* transcript in KO mice was confirmed in mouse cortex, one of the tissues with high *Ctrp10* expression (WT, *n* = 5; KO, *n* = 5). All expression levels were normalized to *β-actin*. All data are presented as mean ± S.E.M. * *P* < 0.05; ** *P* < 0.01; *** *P* < 0.001.

### Generation of *Ctrp10* knockout (KO) mice

We used mice lacking CTRP10 to address whether this secreted protein has a metabolic role in vivo. The mouse *Ctrp10* gene consists of two exons (Fig. 1E). The CRISPR-Cas9 method was used to remove the entire protein coding region spanning exon 1 and 2, thus ensuring a complete null allele (Fig. 1E-F). The targeted allele was confirmed by sequencing. As expected, based on the gene deletion strategy, the *Ctrp10* transcript was absent from KO mice (Fig. 1G).

### CTRP10 is largely dispensable for metabolic homeostasis in young mice fed a control low-fat diet

We first addressed the requirement of CTRP10 in maintaining baseline metabolic homeostasis by assessing metabolic parameters of mice fed a LFD. The body weight and body composition of male mice fed a LFD were not different between genotypes (Fig. 2A-B). By 20 weeks of age, female KO mice fed LFD had a modestly higher body weight relative to WT controls (Fig. 2 C), though the body composition was not different between genotypes (Fig. 2D). Food intake, physical activity, and energy expenditure as measured by indirect calorimetry were also not different between genotypes of either sex across the circadian cycle (light and dark) and metabolic states (*ad libitum* fed, fasted, refed) (Fig. 2E-J). Because *Ctrp10* expression is regulated by nutritional states (Fig. 1C), we assessed serum metabolite levels in WT and KO mice in response to fasting and refeeding. No significant differences in fasting and refeeding blood glucose, serum insulin, triglyceride, cholesterol, non-esterified free fatty acids (NEFA), and β-hydroxybutyrate levels were observed between genotypes of either sex, except the female KO mice had slightly lower fasting β-hydroxybutyrate levels (Fig. 3A-B). We performed glucose and insulin tolerance tests to determine any potential differences in glucose handling capacity and insulin sensitivity. No significant differences in glucose and insulin tolerance were noted between genotypes of either sex (Fig. 3C-F). Together, these data indicate that CTRP10 is dispensable for metabolic homeostasis when mice are young (< 20 weeks old) and fed a LFD.

**Figure 2.**
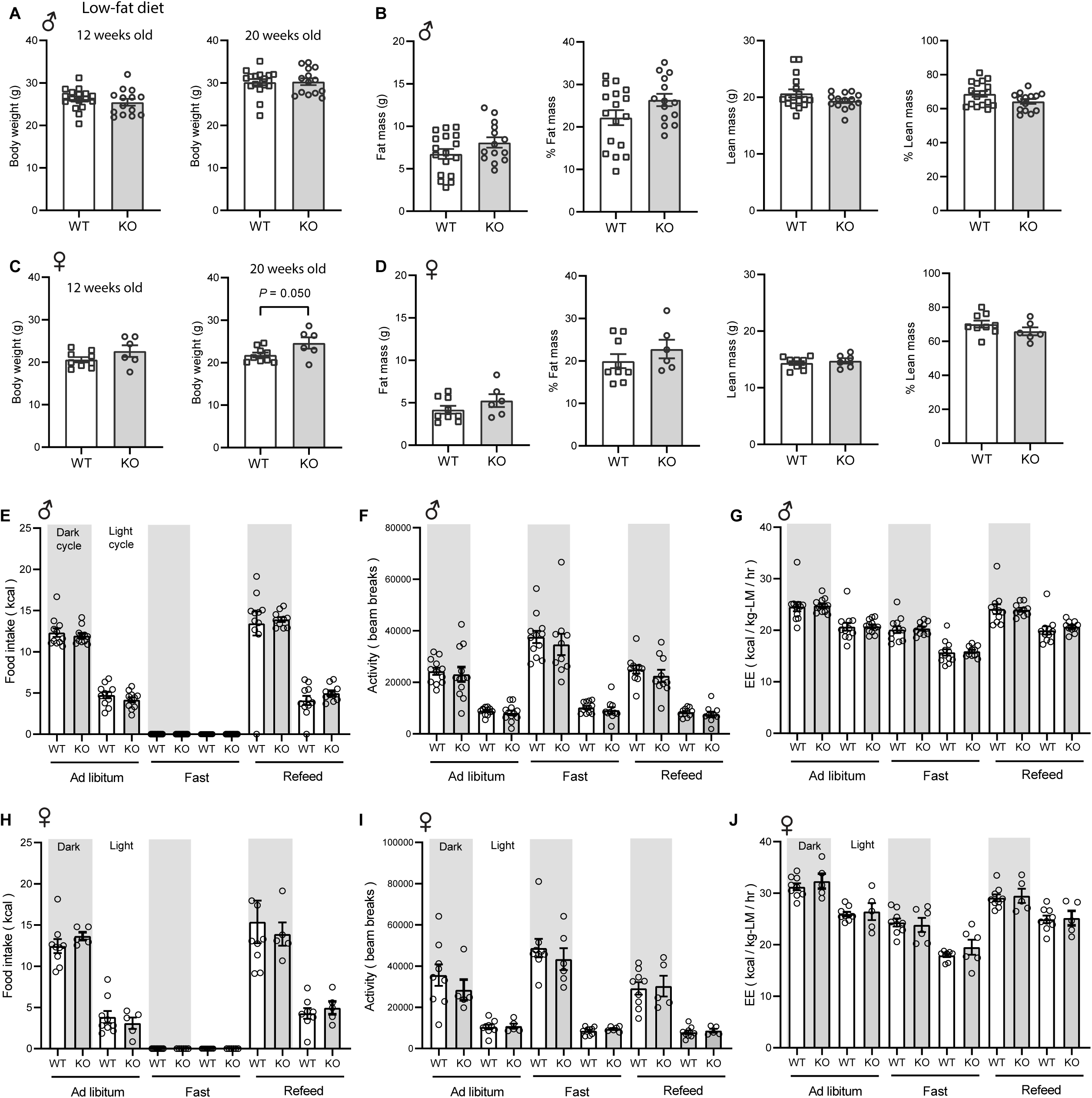
*Ctrp10*-KO mice fed a low-fat diet have normal body weight and energy balance. **(A-B)** Body weight **(A)** and body composition analysis **(B)** of fat mass, % fat mass (relative to body weight), lean mass, and % lean mass of WT (*n* = 17) and KO (*n* = 14) male mice at 18 weeks of age. **(C-D)** Body weight (**C**) and body composition analysis (**D**) of fat mass, % fat mass (relative to body weight), lean mass, and % lean mass of WT (*n* = 9) and KO (*n* = 6) female mice at 13 weeks of age. **(E-G)** Food intake, physical activity, and energy expenditure (EE) in male mice at 18 weeks of age across the circadian cycle (light and dark) and metabolic states (ad libitum fed, fasted, refed) (WT, *n* = 11-12; KO, *n* = 10-12). **(H-J)** Food intake, physical activity, and energy expenditure in female mice at 13 weeks of age (WT, *n* = 9; KO, *n* = 6). All data are presented as mean ± S.E.M.

**Figure 3.**
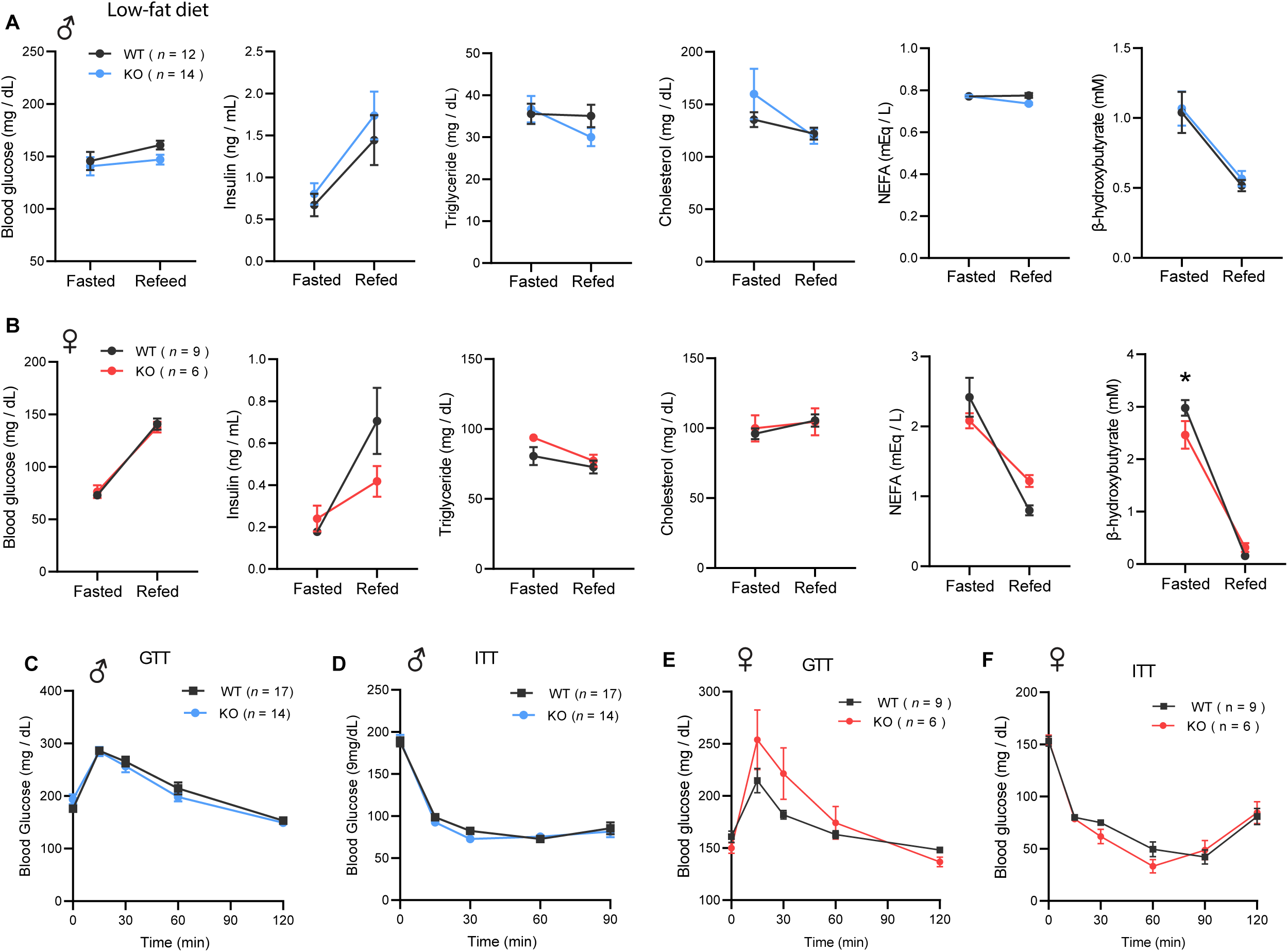
*Ctrp10*-KO mice fed a low-fat diet have normal fasting-refeeding response and glucose homeostasis. (A-B) Overnight fasted and refed blood glucose, serum insulin, triglyceride, cholesterol, non-esterified free fatty acids (NEFA), and β-hydroxybutyrate levels in male **(A)** and female **(B)** mice. **(C-D)** Blood glucose levels during glucose tolerance tests (GTT; **C**) and insulin tolerance tests (ITT; **D**) in WT (*n* = 17) and KO (*n* = 14) male mice at 12 weeks of age. **(E-F)** Blood glucose levels during glucose tolerance tests (GTT; **E**) and insulin tolerance tests (ITT; **F**) in WT (*n* = 9) and KO (*n* = 6) female mice at 20 and 21 weeks of age, respectively. All data are presented as mean ± S.E.M. * *P* < 0.05 (two-way ANOVA with Sidak’s post hoc tests).

### CTRP10-deficient female mice on a low-fat diet develop obesity with age

Because female KO mice were slightly heavier at 20 weeks of age (Fig. 2C), we suspected the weight may diverge further with age. Consequently, we monitored the body weight of female mice fed LFD over an extended period. Indeed, the female KO mice gained significantly more weight and adiposity with age (Fig. 4A-C). By the time the mice reached 40 weeks of age, female KO mice weighed ∼ 6 g (20 %) heavier than the WT controls. Consistent with greater adiposity, the adipocyte cell size (cross-sectional area) was also significantly larger in both gonadal (visceral) white adipose tissue (gWAT) and inguinal (subcutaneous) white adipose tissue (iWAT) (Fig. 4D-E). Increased weight gain over time was not attributed to differences in food intake, as measured manually over a 24 h period (Fig. 4F). Fecal output, frequency, and energy content were also not different between genotypes (Fig. 4G), suggesting that weight gain was not due to greater nutrient absorption. Deep colon temperatures in both light and dark cycle were also not different between genotypes (Fig. 4H). Indirectly calorimetry analyses also revealed no significant differences between genotypes in food intake, physical activity, and energy expenditure across the circadian cycle and metabolic states (*ad libitum* fed, fasted, refed) (Fig. 4I-K). Despite significantly greater body weight and adiposity, female KO mice had the same metabolic profile as the lean WT controls. Given the important role of sex hormone in systemic metabolism, we measured serum estradiol levels and they were not significantly different between WT and KO female mice (Fig. 4 - figure supplement 1). There were no differences in fasting blood glucose, serum insulin, triglyceride, cholesterol, NEFA, and β-hydroxybutyrate levels between genotypes (Fig. 4L). Interestingly, VLDL-TG levels were lower in female KO mice whereas HDL-cholesterol level was not different between genotypes (Fig. 4M). Direct assessments of glucose handling capacity and insulin sensitivity by glucose and insulin tolerance tests, respectively, also revealed no differences between genotypes (Fig. 4N-O). Measurements of mitochondrial respiration in skeletal muscle (gastrocnemius) showed no significant differences in respiration through complex I (CI), CII, and CIV between genotypes; in contrast, *Ctrp10* KO female mice had higher respiration at CIV relative to WT controls (Fig. 4 - figure supplement 2). Together, these data indicate that *Ctrp10*-KO female mice fed LFD develop obesity, but preserve a largely healthy metabolic profile similar to the much leaner WT female mice.

**Figure 4.**
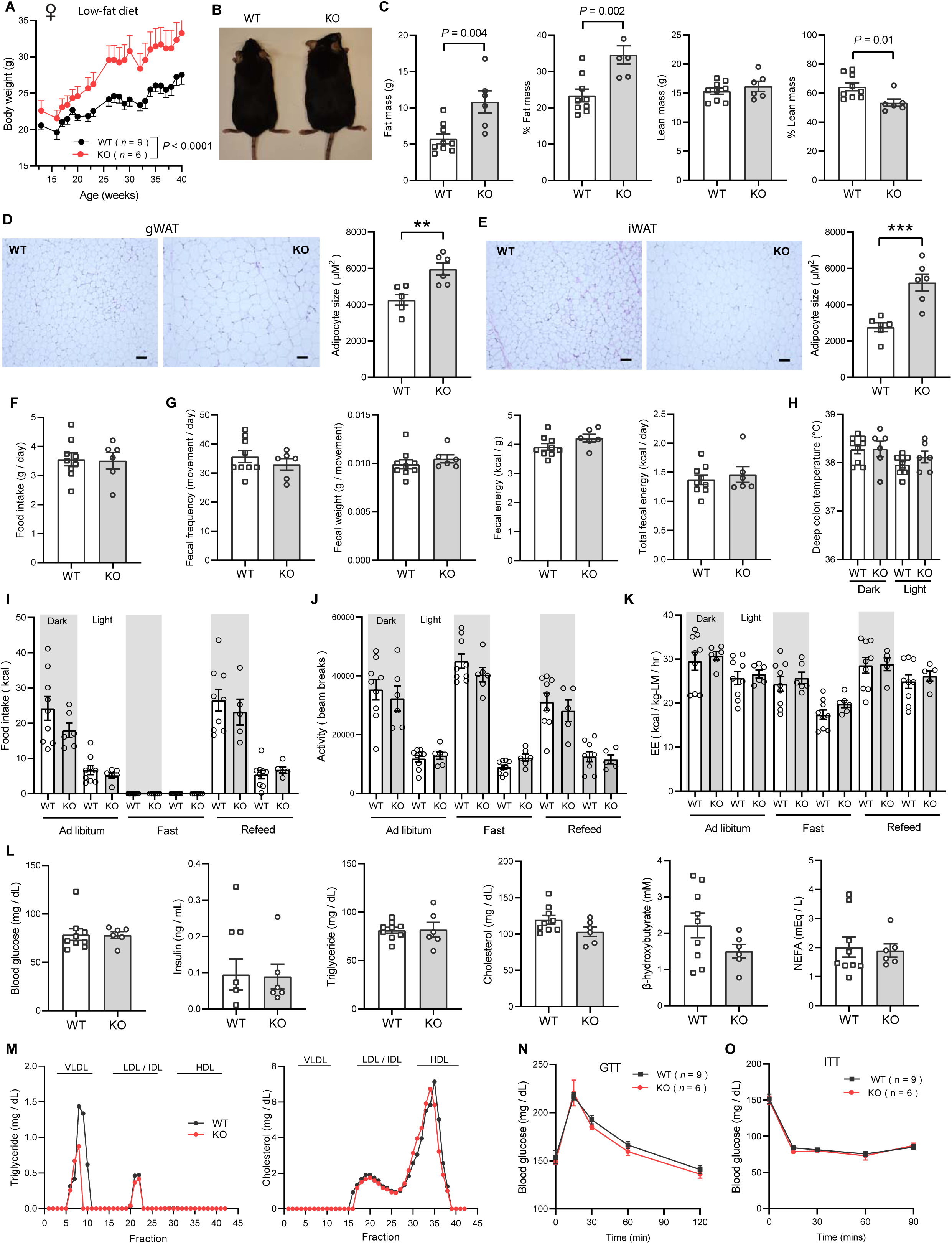
*Ctrp10*-KO female mice on a low-fat diet develop obesity with age. **(A)** Body weights over time of WT and KO female mice fed a low-fat diet (LFD). **(B)** Representative image of WT and KO female on LFD for 40 weeks. **(C)** Body composition analysis of WT (*n* = 9) and KO (*n* = 6) female mice fed a LFD. **(D)** Representative H&E stained histology of gonadal white adipose tissue (gWAT) and the quantification of adipocyte cell size (*n* = 6 per genotype). Scale bar = 100 μM. **(E)** Representative H&E stained histology of inguinal white adipose tissue (iWAT) and the quantification of adipocyte cell size (*n* = 6 per genotype). Scale bar = 100 μM. **(F)** 24-hr food intake data measured manually. **(G)** Fecal frequency, fecal weight, and fecal energy over a 24 hr period. **(H)** Deep colon temperature measured at the light and dark cycle. **(I-K)** Food intake, physical activity, and energy expenditure in female mice across the circadian cycle (light and dark) and metabolic states (ad libitum fed, fasted, refed) (WT, *n* = 9; KO, *n* = 6). Indirect calorimetry analysis was performed after female mice were on LFD for 30 weeks. **(L)** Overnight (16-hr) fasted blood glucose, serum insulin, triglyceride, cholesterol, non-esterified free fatty acids, and β-hydroxybutyrate levels. **(M)** Very-low density lipoprotein-triglyceride (VLDL-TG) and high-density lipoprotein-cholesterol (HDL-cholesterol) analysis by FPLC of pooled (*n* = 6-7 per genotype) mouse sera. **(N)** Blood glucose levels during glucose tolerance tests (GTT). **(O)** Blood glucose levels during insulin tolerance tests (ITT). GTT and ITT were performed when the female mice reached 28 and 29 weeks of age, respectively. WT, *n* = 9; KO, *n* = 6.

### Rapid weight gain in CTRP10-deficient female mice fed a high-fat diet

Next, we challenged the mice with a HFD to determine if the sex-dependent effects on body weight become more pronounced. When fed a HFD, body weight gain and body composition were not different between genotypes in male mice (Fig. 5A-B). Food intake, physical activity, and energy expenditure were also not different between genotypes in male mice across the circadian cycle (light and dark) and metabolic states (*ad libitum* fed, fasted, refed) (Fig. 5C-E). In striking contrast, female KO mice gained weight rapidly on HFD (∼9 g or 28% heavier) and had greater adiposity than the WT controls (Fig. 5F-H). Weight gain in female mice was not due to differences in food intake, body temperature, or changes in fecal output, frequency, and energy content between genotypes (Figure 5 - figure supplement 1). We measured serum estradiol levels and they were not significantly different between WT and KO female mice on HFD (Fig. 5 - figure supplement 2). Metabolic cage analysis also revealed no differences in caloric intake, physical activity, and energy expenditure between genotypes in female mice (Fig. 5I-K). The ANCOVA analysis of energy expenditure using body weight as a covariate also did not reveal any differences between genotypes in female mice (Fig. 5 L). Interestingly, the respiratory quotient (RER) was significantly lower in female KO mice relative to WT controls, especially during fasting and refeeding (Fig. 5M), suggesting a greater reliance on lipid substrates for energy metabolism during those periods. Interestingly, hepatic, but not skeletal muscle, mitochondrial respiration through CII and CIV was lower in *Ctrp10* KO female mice relative to WT controls on HFD (Fig. 5 - figure supplement 3). Together, these data indicate that CTRP10 is required for female-specific body weight control in response to caloric surplus, but neither food intake, physical activity level, nor whole-body energy expenditure could account for the marked increase in body weight and adiposity.

**Figure 5.**
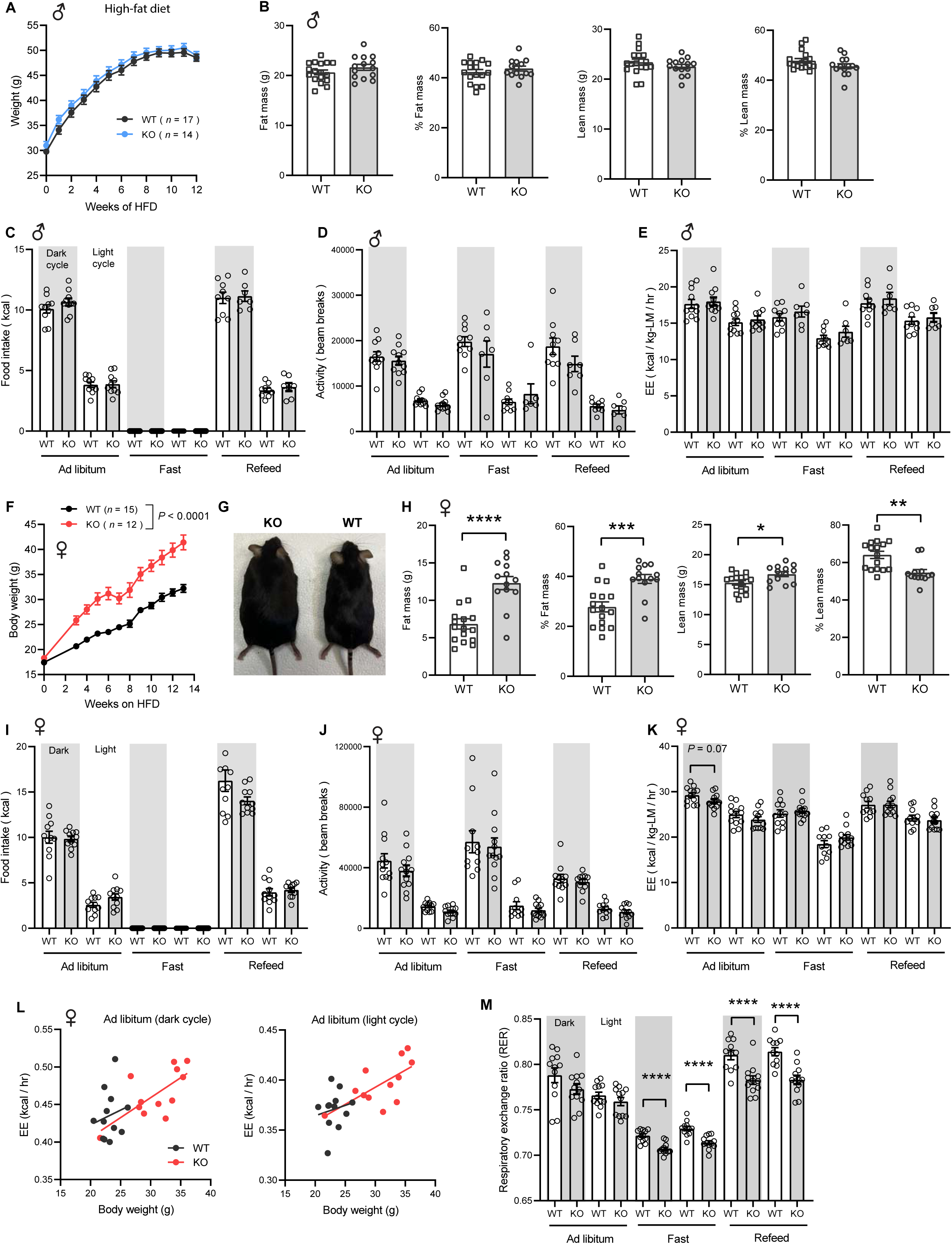
Sexually dimorphic response of *Ctrp10*-KO mice to an obesogenic diet. **(A)** Body weights over time of WT and KO male mice fed a high-fat diet (HFD). **(B)** Body composition analysis of WT (*n* = 17) and KO (*n* = 14) male mice fed a HFD for 9 weeks. **(C-E)** Food intake, physical activity, and energy expenditure in male mice across the circadian cycle (light and dark) and metabolic states (*ad libitum* fed, fasted, refed) (WT, *n* = 11; KO, *n* = 11). Indirect calorimetry analysis was performed after male mice were on HFD for 10 weeks. **(F)** Body weights over time of WT and KO female mice fed a high-fat diet. **(G)** Representative image of WT and KO female mice after 13 weeks of high-fat feeding. **(H)** Body composition analysis of WT (*n* = 17) and KO (*n* = 13) female mice on HFD for 6 weeks. **(I-K)** Food intake, physical activity, and energy expenditure in female mice (WT, *n* = 11-12; KO, *n* = 12) across the circadian cycle (light and dark) and metabolic states (*ad libitum fed*, fasted, refed). Indirect calorimetry analysis was performed after female mice were on HFD for 6 weeks. **(L)** ANCOVA analysis of energy expenditure using body weight as a covariate. **(M)** Respiratory exchange ratio (RER). All data are presented as mean ± S.E.M. * *P* < 0.05; ** *P* < 0.01; *** *P* < 0.001; **** *P* < 0.0001

### Obesity is uncoupled from insulin resistance and dyslipidemia in CTRP10-deficient female mice fed a HFD

To evaluate whether the increased adiposity in KO females led to metabolic dysregulation, we again measured fasting and refeeding responses in WT and *Ctrp10*-KO mice fed a HFD. No differences in fasting and refeeding blood glucose, serum insulin, triglyceride, cholesterol, NEFA, and β-hydroxybutyrate levels were noted between genotypes in male mice (Fig. 6A). Female KO mice, however, had higher fasting blood glucose and serum insulin levels, and lower β-hydroxybutyrate levels compared to the WT controls (Fig. 6B). In the refed state, serum insulin levels continued to be significantly higher in female KO mice. Unlike the WT female mice where refeeding markedly lowered serum β-hydroxybutyrate (ketone) levels as expected, KO female mice appeared unable to suppress serum β-hydroxybutyrate levels in response to refeeding (Fig. 6B).

**Figure 6.**
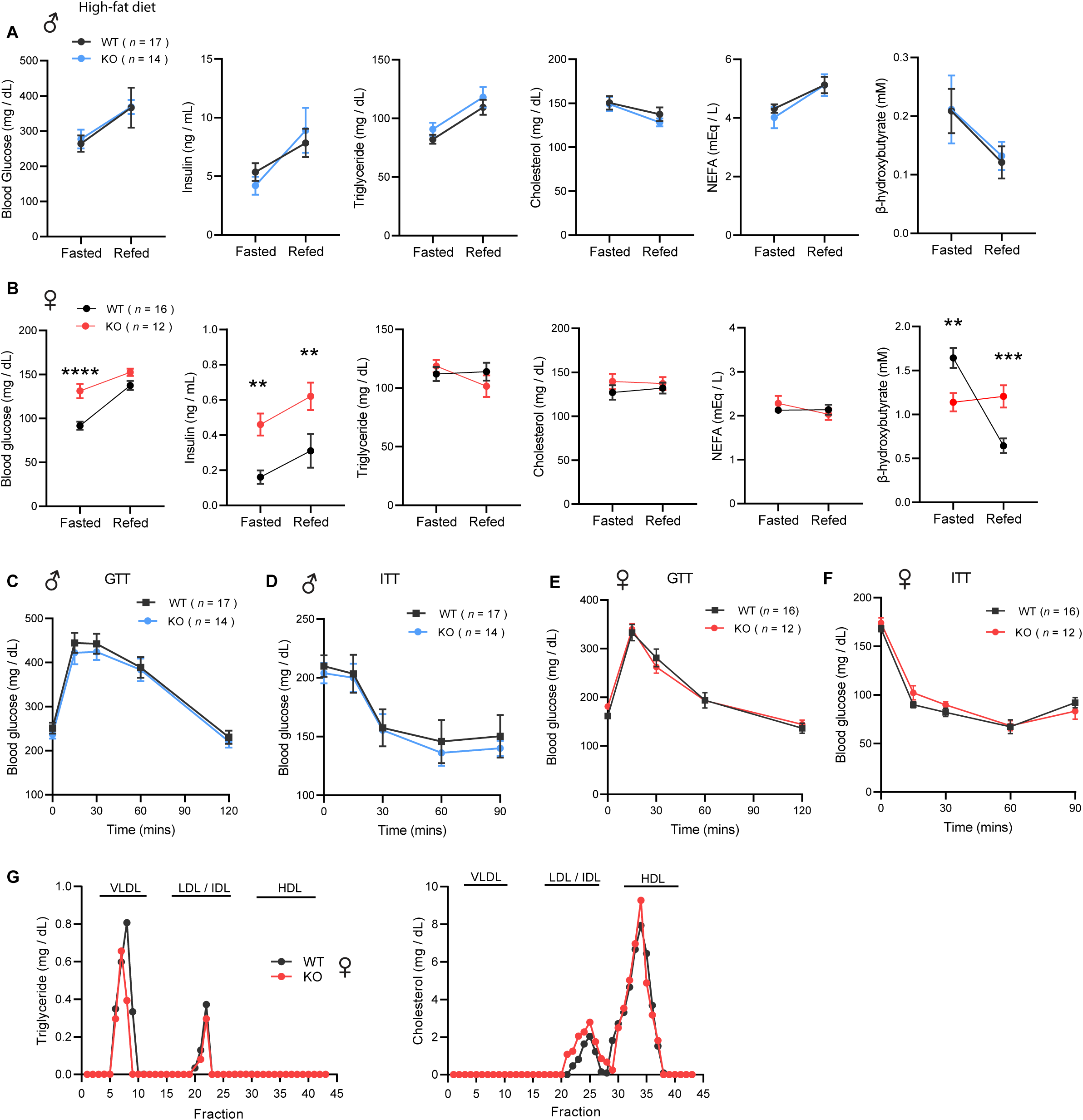
*Ctrp10*-KO mice on a high-fat diet have normal glucose and insulin tolerance. **(A-B)** Overnight fasted and refed blood glucose, serum insulin, triglyceride, cholesterol, non-esterified free fatty acids (NEFA), and β-hydroxybutyrate levels in male **(A)** and female **(B)** mice fed a HFD for 10 weeks. **(C-D)** Blood glucose levels during glucose tolerance tests (GTT; **C**) and insulin tolerance tests (ITT; **D**) in WT (*n* = 17) and KO (*n* = 14) male mice fed a HFD for 10 weeks. **(E-F)** Blood glucose levels during glucose tolerance tests (GTT; **E**) and insulin tolerance tests (ITT; **F**) in WT (*n* = 16) and KO (*n* = 12) female mice fed a HFD for 8 weeks. **(G)** VLDL-TG and HDL-cholesterol analysis by FPLC of pooled female mouse sera. All data are presented as mean ± S.E.M. ** *P* < 0.01; *** *P* < 0.001; **** *P* < 0.0001 (two-way ANOVA with Sidak’s post hoc tests for fasted/refed data).

Consistent with the fasting blood glucose and insulin data, glucose handling capacity and insulin sensitivity assessments by glucose and insulin tolerance in HFD-fed male mice also revealed no differences between genotypes (Fig. 6C-D). Female KO mice, however, had higher fasting blood glucose and serum insulin levels suggesting the presence of mild insulin resistance (Fig. 6B). We therefore expected to see differences in either glucose and/or insulin tolerance tests. To our surprise, the rate of glucose clearance in response to glucose or insulin injection was virtually identical between WT and KO female mice (Fig. 6E-F), suggesting no difference in insulin sensitivity between genotypes. VLDL-TG and HDL-cholesterol profiles were also indistinguishable between WT and KO female mice (Fig. 6G). Altogether, these data indicate that CTRP10 is not required for metabolic homeostasis in male mice challenged with a HFD. In female mice, however, loss of CTRP10 markedly promotes weight gain in the face of caloric surplus, but, paradoxically, the excess adiposity is largely uncoupled from obesity-linked insulin resistance and dysregulated glucose and lipid metabolism.

### Obesity is uncoupled from adipose dysfunction and hepatic steatosis in *Ctrp10*-KO female mice fed a HFD

Consistent with greater fat mass in visceral (gonadal) fat depot of *Ctrp10*-KO female mice (Fig. 7A), histological analysis and quantification also indicated significantly larger adipocyte cell size (Fig. 7B). Although the subcutaneous (inguinal) fat pad weight was also significantly heavier in female KO mice (Fig. 7C), the adipocyte cell size was marginally bigger but not significant (Fig. 7D). A bigger fat pad with only marginally larger cell size suggests greater adipocyte hyperplasia in the subcutaneous depot. Increased adipogenesis in response to caloric surfeit is known to be associated with improved systemic metabolic profile (68).

**Figure 7.**
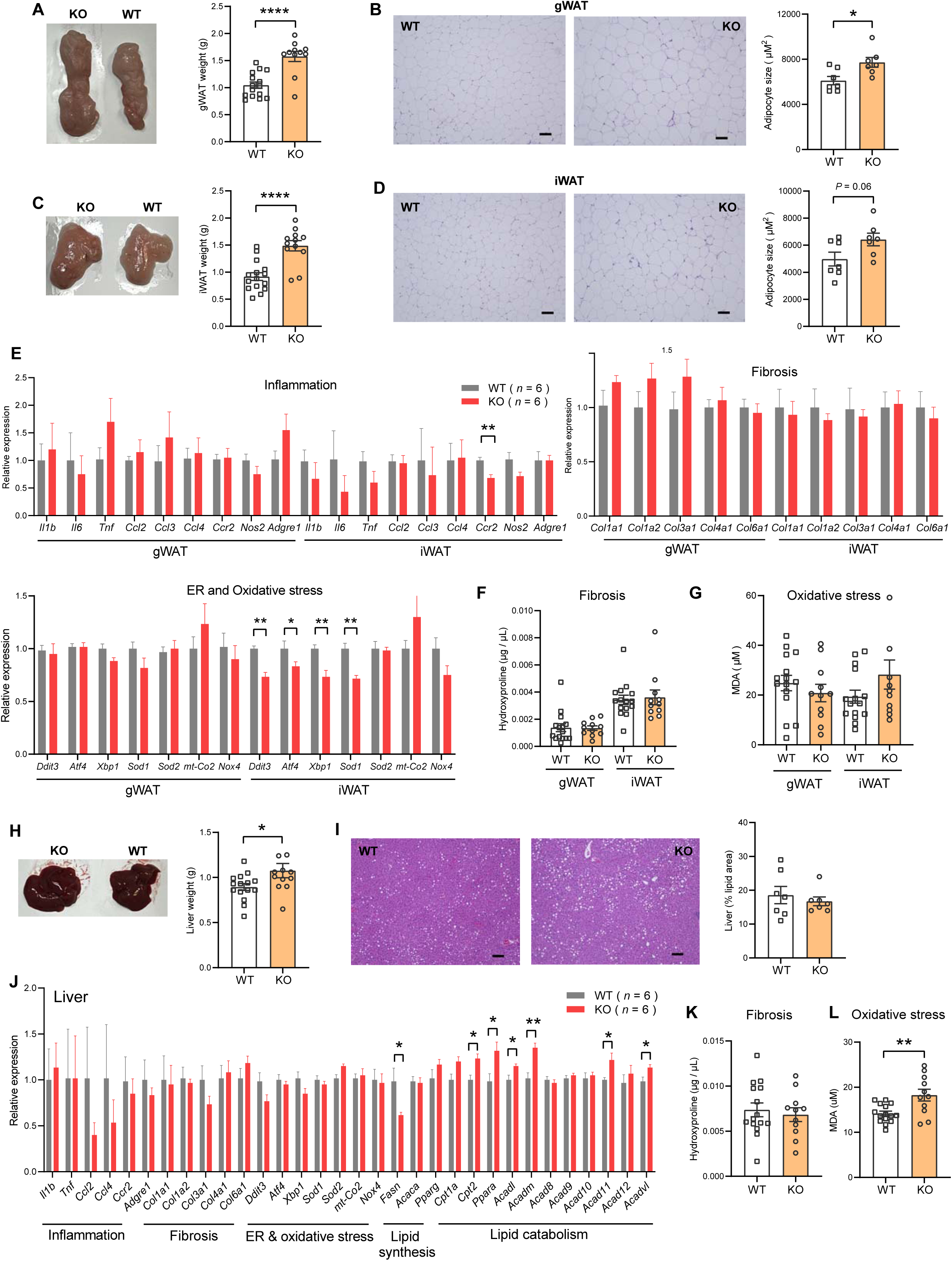
*Ctrp10*-KO female mice fed a HFD do not develop adipose tissue dysfunction and fatty liver. **(A)** Representative images of dissected gonadal white adipose tissue (gWAT) and the quantification of gWAT weight in WT (*n* = 15) and KO (*n* = 12) female mice fed a HFD for 14 weeks. **(B)** Representative H&E stained histological sections of gWAT and the quantification of adipocyte cell size (*n* = 7 per genotype). Scale bar = 100 μM. **(C)** Representative images of dissected inguinal white adipose tissue (iWAT) and the quantification of iWAT weight in WT (*n* = 15) and KO (*n* = 12) female mice. **(D)** Representative H&E stained histological sections of iWAT and the quantification of adipocyte cell size (*n* = 7 per genotype). Scale bar = 100 μM. **(E)** Expression of genes associated with inflammation, fibrosis, ER and oxidative stress in gWAT and iWAT of WT (*n* = 6) and KO (*n* = 6) female mice fed a HFD for 14 weeks. Gene expression data were obtained from RNA-seq. **(F-G)** Quantification of hydroxyproline (marker of fibrosis) and malondialdehyde (MDA; marker of oxidative stress) in gWAT and iWAT. WT, *n* = 15; KO, *n* = 11. **(H)** Representative images of dissected liver and the quantification of liver weight in WT (*n* = 15) and KO (*n* = 12) female mice. **(I)** Representative H&E stained histological sections of liver and the quantification of hepatic lipid content (% lipid area; *n* = 7 per genotype). Scale bar = 100 μM. **(J)** Hepatic expression of genes associated with inflammation, fibrosis, ER and oxidative stress, lipid synthesis, and lipid catabolism in WT and KO female mice. Gene expression data were obtained from RNA-seq. **(K-L)** Quantification of hydroxyproline (marker of fibrosis) and malondialdehyde (MDA; marker of oxidative stress) in liver. WT, *n* = 15; KO, *n* = 11. All data are presented as mean ± S.E.M. * *P* < 0.05; ** *P* < 0.01.

Obesity is known to be associated with low-grade inflammation (69), fibrosis (70), and ER and oxidative stress (71, 72). Despite marked differences in body weight and adiposity, the expression of genes associated with inflammation (except for *Ccr2*), fibrosis, oxidative stress in gWAT and iWAT were not significantly different between female KO mice and WT controls (Fig. 7E). The expression of some genes associated with ER stress (e.g., *Ddit3*/*CHOP*, *Atf4*, *Xbp1*) in iWAT were actually lower in female KO mice (Fig. 7E). Corroborating the gene expression data, quantification of hydroxyproline (marker of fibrosis) and malondialdehyde (marker of oxidative stress) revealed no significant differences between genotypes (Fig. 7F-G).

The liver weight of female KO mice was modestly increased (Fig. 7F), but when normalized to body weight it was not significantly different from WT controls (2.76 % in WT and 2.60% in KO, *P* = 0.24). Histological analysis and quantification revealed no differences in hepatic lipid content (% lipid area) between genotypes (Fig. 7G). Interestingly, although hepatic fat content was similar between genotypes, the expression of lipogenic genes (e.g., *Fasn*) was lower and fat catabolism genes (e.g., *Cpt2*, *Ppara*, *Acadl*, *Acadm*, *Acad11*, *Acadvl*) was higher in female KO mice (Fig. 7H). The expression of genes associated with inflammation, fibrosis, ER and oxidative stress in liver were not significantly different between genotypes (Fig. 7H). Consistent with the gene expression data, quantification of hydroxyproline (marker of fibrosis) in the liver revealed no significant difference between genotypes (Fig. 7K). The *Ctrp10* KO female mice, however, had higher levels of malondialdehyde (a marker of oxidative stress) in the liver, suggesting a modest increase in oxidative stress (Fig. 7L). Altogether, these data indicate that obesity is largely uncoupled from inflammation, fibrosis, ER and oxidative stress in *Ctrp10* KO female mice.

### Transcriptomic and pathway changes associated with the metabolically healthy obesity phenotype in *Ctrp10* KO female mice

To define the specific mechanisms mediating the female-specific effects of *Ctrp10* ablation on favorable metabolic outcomes, four major metabolic tissues (gWAT, iWAT, liver, skeletal muscle) from female WT and KO mice fed a HFD were subjected to RNA-sequencing (Figure 8 - Source data 1-8). Comparison of differentially expressed genes (DEGs) via limma (73) showed robust changes across tissues, with the largest changes seen in the liver (Fig. 8A-D). In liver, gWAT, and muscle, we observed comparable numbers of DEGs that were up- and down-regulated, whereas more genes were transcriptionally suppressed in the iWAT of *Ctrp10* KO female mice (Fig. 8E, top panel). While significant DEGs were identified in all 4 tissues, only limited overlap was observed between the DEGs in each tissue (Fig. 8E, bottom panel). Gene set enrichment analyses of the DEGs highlighted distinct and shared processes up- or down-regulated across the four tissues (Fig. 8F). Pathways and processes related to lipid metabolism and estrogen receptor were the top-ranked up-regulated enrichments across tissues (Fig. 8F, top panel), whereas processes related to blood clotting and lipoprotein metabolism were the top-ranked down-regulated enrichments (Fig. 8F, bottom panel).

**Figure 8.**
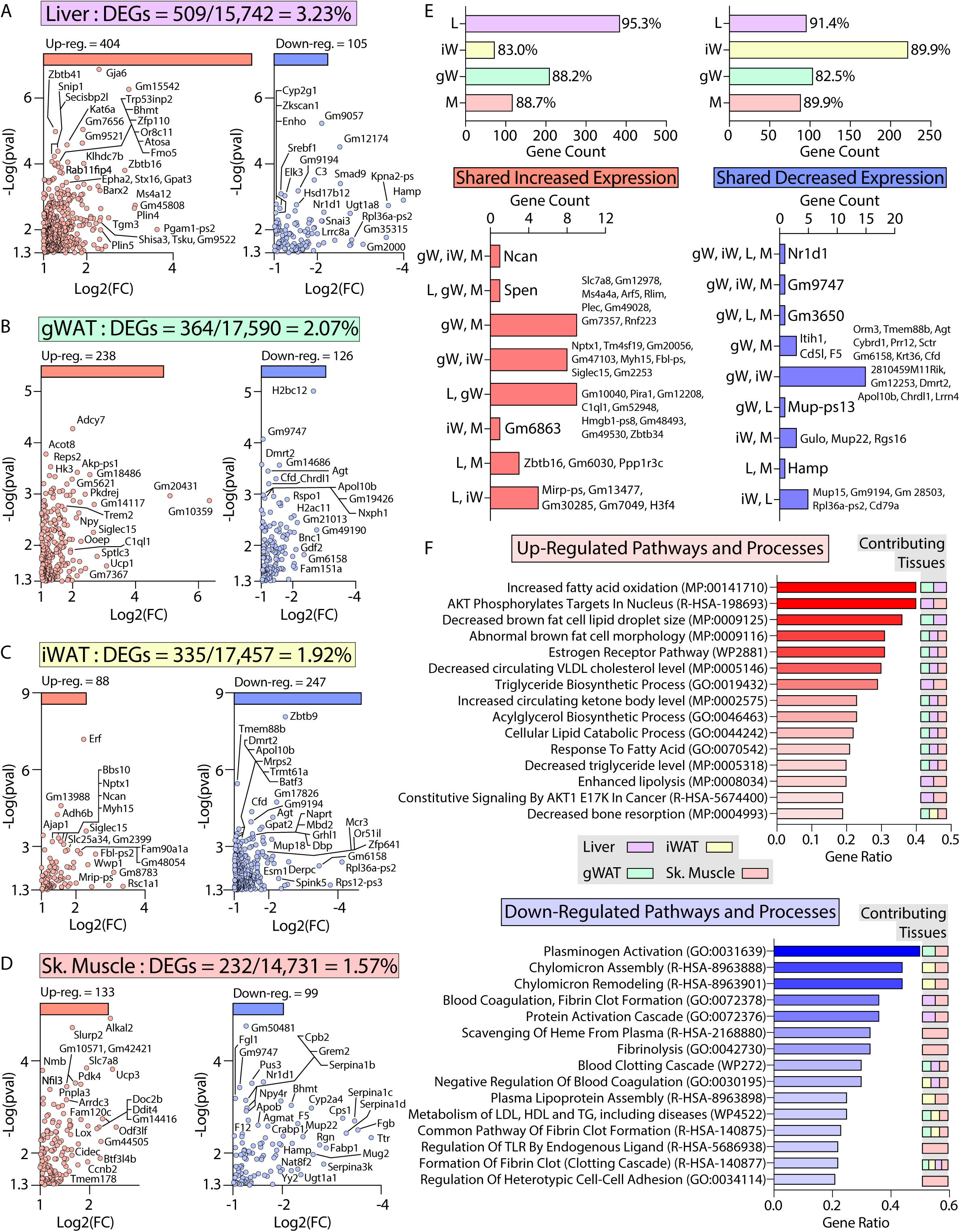
Transcriptomic analysis of liver, adipose tissue, and skeletal muscle of female *Ctrp10* KO mice fed a high-fat diet. **(A-D)** Cropped volcano plot views of all differentially expressed genes (DEGs, Log2(Fold Change) >1 or <-1 with a *p*-value <0.05) of the liver, gonadal white adipose tissue (gWAT), inguinal WAT (iWAT), or skeletal muscle (gastrocnemius). **(E)** Overlap analysis of tissue DEGs showing (top panel) expression unique to gonadal white adipose tissue (gW), inguinal white adipose tissue (iW), liver (L), or skeletal muscle (M). Percent (%) represents percent DEGs unique to each tissue. Bottom panel show DEGs shared across multiple tissues, with all the shared DEGs listed. **(F)** Enrichr analysis (134) of biological pathways and processes significantly (p<0.01) affected across the CTRP10 deficient female mice. Top pathways and processes derived from Gene Ontology (GO), Reactome (R-HAS), WikiPathway human (WP), and mammalian phenotype (MP). All up- or down-regulated DEGs across all tissues were used for analysis. The tissues contributing to the highest ranked pathways and processes are specified. *n* = 6 KO and 6 WT for RNA-seq experiments.

Of the DEGs, we found significant changes across tissues in relevant classes of genes that encode proteins involved in gene expression (e.g., transcription factors), signaling (e.g., receptors), tissue crosstalk (e.g., secreted proteins), and metabolism (Fig. 9). Notably, the nuclear receptor, *Nr1d1* (also known as *Rev-Erbα*), is the only gene consistently suppressed across all four tissues (liver, gWAT, iWAT, and muscle) of *Ctrp10* KO female mice (Fig. 8E lower panel and Fig. 9). Interestingly, global deletion of *Nr1d1* promotes lipogenesis, adipose tissue expansion, and obesity (74, 75). Although the whole-body and adipose-specific *Nr1d1* KO mice fed with HFD become markedly obese, the obesity is not accompanied by insulin resistance, adipose tissue inflammation and fibrosis (75, 76). Like the *Ctrp10* KO female mice, HFD-fed mice lacking NR1D1 can maintain a relatively healthy metabolic profile despite being strikingly obese. Since only male mice were used in these previous studies, we do not know whether female mice lacking NR1D1 would also exhibit similar insulin-sensitive obesity phenotype. In WT mice, NR1D1 acts as a transcriptional repressor of metabolic genes whose expression are upregulated by high-fat feeding; loss of NR1D1 is thought to result in the de-repression of these genes, leading to greater lipid synthesis and fat mass accrual in response to caloric excess (76). Thus, the suppression of *Nr1d1* expression—mimicking NR1D1 deficiency— across tissues in HFD-fed *Ctrp10* KO female mice may contribute to benign fat mass expansion without the accompanying adipose tissue fibrosis, inflammation, and oxidative stress.

**Figure 9.**
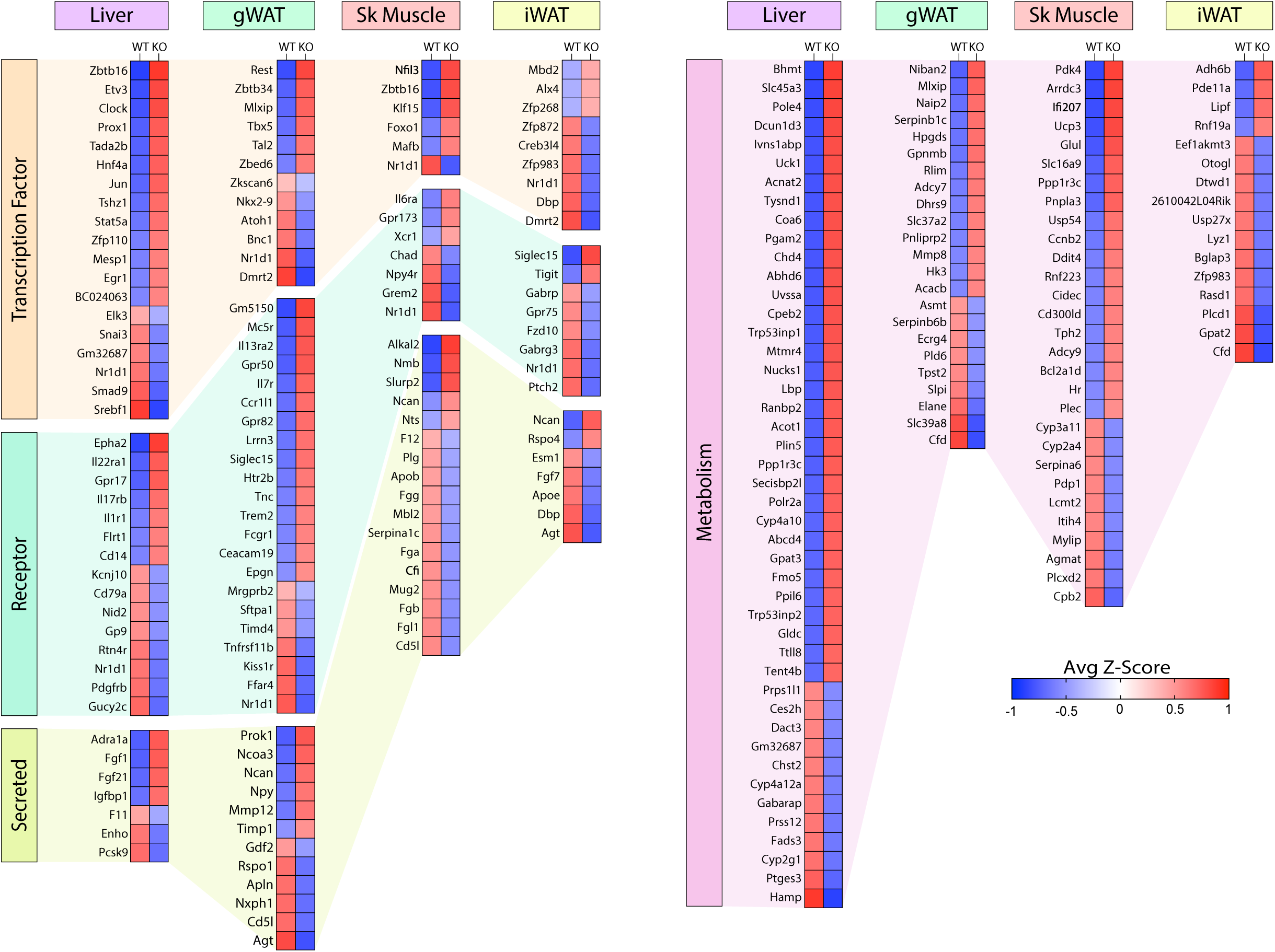
Loss of CTRP10 induces significant and wide-spread alterations in the expression of key transcription factors, secreted protein, membrane receptors, and metabolism-associated genes. **(A-D)** Selected genes from the DEG list of each tissue organized based on gene type (genes encoding transcription factors, secreted proteins, receptors, and proteins involved in metabolism) and ranked from highest to lowest average row z-score. Z-score is defined as z = (x-μ)/σ, where x is the raw score (gene transcript level), μ is the population mean (i.e., mean of gene expression across both WT and KO samples), and σ is the population standard deviation. *n* = 6 per genotype.

Among the upregulated genes, *Fgf21* and *Fgf1* were significantly elevated in the liver of *Ctrp10* KO female mice (Fig. 9A). FGF21 is an hepatokine known to improve systemic insulin sensitivity and to promote a favorable metabolic profile in diet-induced obese mice (77). While hepatic *Fgf21* expression is significantly elevated, its systemic circulating levels were only found to be higher (*P* = 0.051) in LFD-fed, but not HFD-fed, KO female mice (Fig. 9 – figure supplement 1); this suggests liver-derived FGF21 may be retained and act locally within the liver parenchyma. FGF1 is another secreted protein that has been shown to dampen hepatic glucose output by suppressing adipose lipolysis (78), improve systemic insulin sensitivity by reducing adipose inflammation (79), and alleviate hepatic steatosis, inflammation, and insulin resistance (80). Thus, upregulated expression of *Fgf21* and *Fgf1* in *Ctrp10* KO female mice could contribute to the MHO phenotype. In addition, the upregulated hepatic expression of IL-22 receptor (*Il22ra1*) in *Ctrp10* KO female mice (Fig. 9A) may confer protection against obesity-associated fatty liver, inflammation and fibrosis (81–83). Further, a marked increase in uncoupling protein 3 (*Ucp3*) and Krüppel-like factor 15 (*Klf15*) expression in the skeletal muscle (Fig. 9C) may promote lipid utilization and help mitigate lipid-induced insulin resistance in *Ctrp10* KO female mice (84, 85). Taken together, these combined changes—at the level of gene expression and biological pathways and processes across tissues—acting in concert likely contribute to the apparently healthy obesity phenotype seen in the KO female mice.

### Conservation of mouse DEG co-correlation in humans highlights sex-specific gene connectivity

Next, we asked whether the female-specific transcriptomic effects across tissues were conserved in humans. To address this, we analyzed transcriptional co-correlation of mouse DEG (Fig. 10) orthologues in GTEx (86), consisting of 210 males and 100 females filtered for comparison of gene expression across tissues (87, 88). Hierarchical clustering of transcriptional correlation of the orthologous DEGs among 4 metabolic tissues— subcutaneous and visceral white adipose tissue, liver, and skeletal muscle—showed differing patterns of gene connectivity between females (Fig. 10A) and males (Fig. 10B). When grouped according to sex in each tissue, the degree of sex-specific gene correlation pairs of DEGs orthologues showed the most significant differences in subcutaneous adipose tissue (Fig. 10C). Given the whole-body metabolic effects of *Ctrp10* ablation in mice, we further examined the degree of sex-dependent DEG co-correlation across metabolic tissues. This analysis showed that human orthologue genes in subcutaneous adipose tissue (Fig. 10D, top row) and liver (Fig. 10D, third row) also exhibited highly significant sex differences in their transcriptional correlation with other DEGs across key metabolic tissues (Fig. 10D). These analyses highlight the sex-specificity of CTRP10 DEG orthologues in humans, suggest possible sex-biased mechanisms of tissue crosstalk, and overall underscores the conservation of the sex-dependent metabolic function of CTRP10.

**Figure 10.**
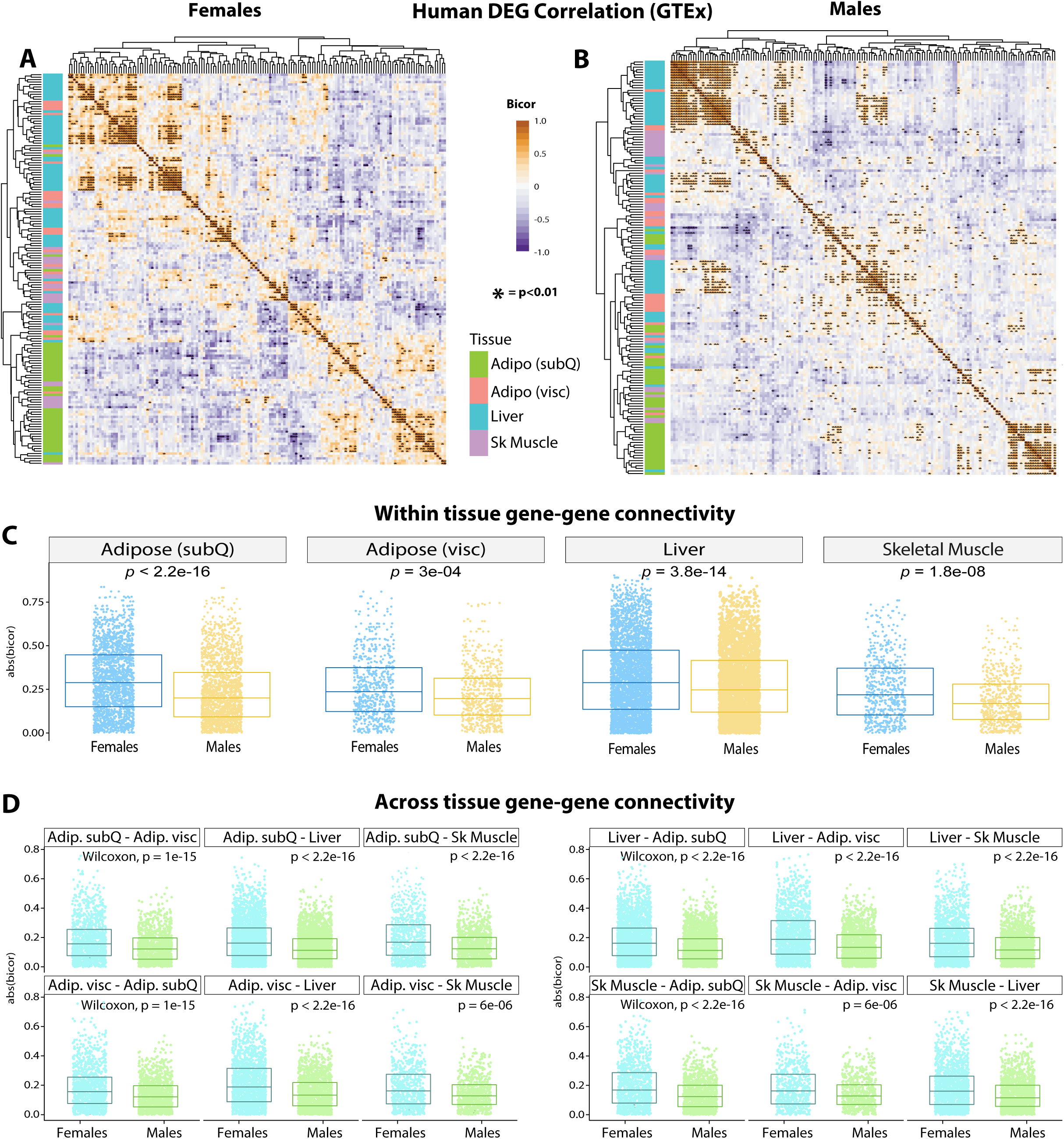
GTEx genetic co-correlation of mouse differentially expressed gene (DEG) orthologues. **(A-B)** Heatmaps showing biweight midcorrelation (bicor) coefficient among human tissue DEG orthologues in females **(A)** and males **(B)** in GTEx. Y-axis color indicates tissue of origin, *P*-value based on students’ regression *P*-value. **(C)** T-tests between correlation coefficient in males and females among all DEG orthologue gene pairs for subcutaneous (SubQ) adipose tissue, visceral (visc) adipose tissue, liver, and skeletal muscle. **(D)** the same as in C, except comparisons are shown for all gene-gene pairs between tissues. For example, the top left graph compared the connectivity of males (blue color) vs females (green color) for correlation between subcutaneous (SubQ) and visceral (Visc) adipose tissue DEG orthologues.

## DISCUSSION

Our current study has established a novel function for CTRP10 in modulating body weight in a sex-specific manner. When mice were fed a control LFD, female *Ctrp10*-KO mice developed obesity with age; increased adiposity, however, did not impair insulin action and glucose and lipid metabolism. When challenged with an obesogenic diet, female *Ctrp10*-KO mice gained weight rapidly. Despite having strikingly higher adiposity and weighing ∼10-11 g (∼28%) more, female KO mice fed a HFD exhibited a metabolic profile largely indistinguishable from the much leaner WT controls. Although female KO mice had higher fasting glucose and insulin levels, direct assessments of glucose metabolism and insulin sensitivity by glucose and insulin tolerance tests, however, revealed no differences between genotypes. Except having lower fasting ketone (β-hydroxybutyrate) levels, the fasting lipid profile, as well as VLDL-TG and HDL-cholesterol levels, of *Ctrp10-*KO female mice resembled the WT controls. The hepatic fat content was also comparable between genotypes. Global transcriptomic profiling across different fat depots and liver did not reveal gene expression signatures associated with elevated inflammation, fibrosis, and ER and oxidative stress. Altogether, these findings suggest that CTRP10 deficiency promotes obesity in females but it also uncouples obesity from insulin resistance, dyslipidemia, steatosis, inflammation, and oxidative stress. Thus, *Ctrp10*-KO female mice represent a novel model of female obesity with largely preserved insulin sensitivity and metabolic health.

Our findings help inform ongoing studies on metabolically healthy obese (MHO) humans (89–94). Because the criteria used to define MHO differs between studies, there is an ongoing debate regarding the prevalence of MHO and what fraction of the MHO population is insulin-sensitive and metabolically healthy (95, 96). Nevertheless, among the obese individuals, there clearly exists a subgroup that maintains long-term normal insulin sensitivity and does not appear to develop any component of the metabolic syndrome (95). MHO is observed in both sexes, but more common in females (6). The underlying mechanism(s) that uncouple obesity from adverse metabolic health in MHO is not well understood (7, 91, 95). It is currently unknown whether males and females with MHO use similar or distinct mechanism to maintain insulin sensitivity and metabolic health. Our findings in *Ctrp10*-KO female, but not male, mice suggest that there may be female-biased mechanism that prevents metabolic deterioration in the face of obesity, thus underscoring the utility of the *Ctrp10*-KO mice as a female mouse model of MHO.

Obesity is frequently associated with insulin resistance, dyslipidemia, fatty liver, oxidative stress, and chronic low-grade inflammation (69, 97, 98). The mechanisms that link obesity to metabolic dysfunctions are complex and multifactorial. There are limited number of mouse models described where obesity is uncoupled from insulin resistance and metabolic health (99–102); in some studies, however, only male mice were used or that the sex of the animals was not specified. In the case of aP2/FABP4 KO male mice, the uncoupling of obesity from insulin resistance was attributed to a marked decrease in TNF*-α* expression in adipose tissue (100). In the case of adiponectin overexpression in leptin-deficient (*ob*/*ob*) male and female mice, a dramatic expansion of the subcutaneous fat pad is thought to promote lipid sequestration in adipose compartment, thus preventing ectopic lipid deposition in non-adipose tissues (e.g., liver, pancreas, muscle) that would otherwise induce insulin resistance (99). Massive obesity with preserved insulin sensitivity is also observed in leptin-deficient (*ob*/*ob*) male and female mice overexpressing the mitochondrial membrane protein, mitoNEET (102). The benign obesity is attributed to the inhibition of iron transport into mitochondria by mitoNEET, leading to reduced mitochondrial activity, fatty acid oxidation, and oxidative stress (102). In the Brd2 hypomorphic mice, severe obesity with lower blood glucose and enhanced glucose tolerance is due to a combination of hyperinsulinemia and marked reduction in macrophage infiltration into fat depot (103). Lastly, in male mice fed a high starch diet, the uncoupling of obesity from insulin resistance is associated with lower ceramide levels in liver and skeletal muscle (101). In all these cases, the uncoupling of obesity from metabolic dysfunction is seen in either male mice only (female mice were not included) or both sexes. These previous studies suggest that multiple mechanisms, not mutually exclusive, can contribute to the MHO phenotype in different mouse models.

In our study, loss of CTRP10 in female mice largely uncoupled obesity from insulin resistance, dyslipidemia, steatosis, inflammation, and ER and oxidative stress. The preservation of insulin sensitivity in *Ctrp10* KO female mice is due, at least in part, to the absence of obesity-linked adipose and liver inflammation, fibrosis, and oxidative stress. These phenotypes typically associated with a favorable metabolic profile are also observed in MHO individuals (90, 104). A healthy adaptive remodeling of white adipose tissues in response to caloric surfeit helps preserve the storage and secretory function of adipocytes (105). The expansion of benign adipose tissues further serves to sequester circulating lipids and prevent their ectopic deposition in non-adipose tissue (e.g., liver and skeletal muscle) which can impair insulin action (106, 107). The MHO phenotype seen in *Ctrp10* KO female mice reinforce the notion that adipose tissue health, rather than abundance, is an important determinant of metabolic health in obesity.

Because lipidomic analysis was not performed—a limitation of this study—we do not know whether *Ctrp10*- KO female mice have reduced ceramide or diacylglycerol levels in liver and skeletal muscle, two lipid species known to antagonize insulin action (108, 109). However, our global transcriptomic and pathway enrichment analysis across visceral and subcutaneous fat depots, liver, and skeletal muscle highlighted the relevant up- and down-regulated pathways and processes (e.g., lipid and lipoprotein metabolism, signaling) that may contribute to the MHO phenotype in *Ctrp10*-KO female mice. How these changes across tissues help to suppress the deleterious effects of obesity and maintain an apparently healthy metabolic profile in *Ctrp10*-KO female mice remains to be fully elucidated. Part of the mechanism may be attributable to the suppression of *Nr1d1* and the upregulated expression of *Fgf1*, *Fgf21*, *Il22ra1*, *Ucp3*, *Klf15*. Increased expression of these genes is known to reduce obesity-linked inflammation, oxidative stress, steatosis, and insulin resistance. Interestingly, global deletion of *Nr1d1* preserves insulin sensitivity despite promoting lipogenesis, adipose tissue expansion, and obesity (74, 75). NR1D1 (also known as REV-ERBα) is a nuclear receptor that acts as a transcriptional repressor (110, 111). Circadian and metabolic genes that normally repressed by NR1D1 are de-repressed in mice lacking NR1D1 globally or in a fat-specific manner, thus leading to fat mass accrual (74, 76). It remains to be determined whether the suppression of *Nr1d1* expression in adipose tissue, liver, and skeletal muscle is indeed responsible, at least in part, for the MHO phenotype seen in *Ctrp10* female mice.

In many single-gene KO mouse models where both sexes are examined, it is often the males that show a more pronounced metabolic phenotype. It is known that C57BL/6J female mice generally gain significantly less weight on HFD compared to male mice (112). Therefore, it is intriguing that *Ctrp10*-KO female mice became obese on a control LFD and gained weight rapidly when fed an obesogenic diet. After twelve weeks on HFD, the body weight of female KO mice was approaching that of WT male mice fed the same diet. What mechanism underlies the sexually dimorphic requirement of CTRP10 for body weight control? We know that the obesity phenotype was not attributed to differences in food intake, physical activity level, body temperature, and energy expenditure between WT and KO female mice. We assume that the methods used to quantify these physiologic parameters are sensitive enough to detect small differences that can give rise to divergent body weight over time. Quantification of fecal output and fecal energy content also revealed no differences between genotypes. Thus, loss of CTRP10 did not affect macronutrient intake and absorption. Although we cannot fully rule out the CNS function of CTRP10/C1QL2 (62), our data do not support a central role for CTRP10 in modulating food intake behavior, locomotor activity, and energy expenditure that affect body weight in female mice.

It is known that reduced estrogen level by ovariectomy or blocking estrogen action in estrogen receptor (ERα) KO mice will cause obesity and metabolic dysfunction in female mice fed a HFD (113–115). Conversely, estradiol supplementation decreases HFD-induced weight gain and improves glucose tolerance and insulin sensitivity (116–118). Estrogen also has the effect of reducing food intake, and promoting physical activity and energy expenditure (119–123). In our study, loss of CTRP10 promotes obesity without altering food intake, physical activity, and energy expenditure. While estrogen’s role cannot be completely ruled out, the fact that estrogen levels were not different between genotypes and that female *Ctrp10*-KO mice developed obesity with largely preserved metabolic health suggest that factors other than altered estrogen level contribute to the insulin-sensitive obesity phenotype. Future studies are warranted to uncover what factor(s) is causally contributing to obesity in female mice lacking CTRP10.

The sex-dependent effects of CTRP10 on metabolism and tissue transcriptomes appear to be conserved in humans. When the human orthologues of the mouse DEGs were used to interrogate the GTEx data, clear patterns of gene connectivity within and across metabolic tissues in females and males were observed, with the strongest sex-specific gene correlations seen in subcutaneous adipose tissue and liver. These findings provide further evidence that CTRP10 modulates tissue transcriptome in a sex-dependent manner. Further, our analyses of sex-dependent DEG co-correlation across metabolic tissues also suggest possible sex-biased mechanisms of inter-organ metabolic signaling between adipose tissue and liver.

CTRP10 has been previously shown to bind to the adhesion GPCR, brain angiogenesis inhibitor-3 (Bai3/Adgrb3) (124). Bai3 is expressed in the brain and peripheral tissues, and it is a promiscuous GPCR that can bind to multiple ligands. In addition to CTRP10 (C1QL2), Bai3 also binds to CTRP11 (C1QL4), CTRP13 (C1QL3), CTRP14 (C1QL1), neuronal pentraxins, and reticulon 4 (RTN4) receptor (56, 58, 59, 124–126). A constitutive, whole-body KO of Bai3 mouse models have recently been generated (127, 128). Both male and female *Bai3* KO mice fed a standard chow have significantly lower body weight, beginning at weaning (3 weeks old) and continue into adulthood (127, 128). Lower body weight in *Bai3* KO mice of either sex is attributed to a reduction in both lean and fat mass, and is associated with higher energy expenditure and reduced food intake in male mice (127). The impact of Bai3 deficiency on systemic metabolism in response to a high-fat diet was not examined. The *Ctrp10* KO mice do not phenocopy the phenotypes of the *Bai3* KO mice. When fed a control low-fat diet, the body weight, food intake, and energy expenditure of *Ctrp10* KO male mice were indistinguishable from WT controls. In striking contrast to *Bai3* KO mice, female *Ctrp10* KO mice fed a low-fat diet began to gain more weight around 20 weeks of age, and by 40 weeks had become visibly obese. While we do not rule out CTRP10-Bai3 signaling axis in modulating energy metabolism in peripheral tissues, our findings in *Ctrp10* KO mice suggest that future works are needed to establish the molecular mechanisms that mediate the systemic metabolic function of CTRP10.

Several limitations of our current study are noted. The lack of reliable antibody specific for CTRP10 preclude the assessment of protein abundance in response to altered nutritional and metabolic states. We used a constitutive whole-body KO mouse model of CTRP10 to interrogate its function. It is unknown whether CTRP10 has a role during development that may influence sex-dependent postnatal weight gain with age or in response to a high-caloric diet. Future studies using conditional KO of *Ctrp10* gene in adult mice can help address this issue. We also did not perform tracer studies to address whether there are anabolic and catabolic changes in glucose and fat metabolism in the liver and adipose tissue of *Ctrp10* KO female mice, as this information may shed light on how different tissues handle macronutrients in the absence of CTRP10. In KO female mice, we observed upregulated expression of genes (e.g., *Fgf1*, *Fgf21*, *Il22ra1*, *Ucp3*, and *Klf15*) in liver or skeletal muscle that potentially contributes to the MHO phenotype. The mechanism by which CTRP10 suppresses—either directly or indirectly—the expression of these genes and their de-repression in the absence of CTRP10 is presently unknown. Although the lack of differences in food intake, physical activity, and energy expenditure between genotypes do not support a central role of CTRP10 in mediating the metabolic phenotypes of *Ctrp10*-KO female mice, a brain-specific KO of *Ctrp10* gene is needed to definitively rule this out. Our phenotypic analyses in the context of metabolism are relatively comprehensive but not exhaustive. Although the metabolic profile of obese *Ctrp10*-KO female mice was largely indistinguishable from the much leaner WT controls, some related aspect of metabolic health (e.g., blood pressure and heart function) may be altered in the absence of CTRP10 which we did not examine.

In summary, we have established the physiologic role and requirement of CTRP10 in modulating body weight in a female-specific manner. Importantly, loss of CTRP10 uncouples obesity from insulin resistance and metabolic dysfunction. The CTRP10-deficient female mice represent a unique and valuable model for dissecting female-biased mechanisms that help preserve metabolic health in the face of positive energy balance and increased adiposity.

## MATERIALS AND METHODS

### Mouse models

Eight-week-old mouse tissues (gonadal and inguinal white adipose tissues, interscapular brown adipose tissue, liver, heart, skeletal muscle, kidney, pancreas, cerebellum, cortex, hippocampus, hindbrain, and hypothalamus) from C57BL/6J male mice (The Jackson Laboratory, Bar Harbor, ME) were collected from fasted and refed experiments as we have previously described (129). For the fasted group (∼10 weeks old), food was removed for 16 h (beginning 10 h into the light cycle), and mice were euthanized 2 h into the light cycle. For the refed group (∼10 weeks old), mice were fasted for 16 h and refed with chow pellets for 2 h before being euthanized. Tissues (white and brown adipose tissues, liver, whole brain, kidney, spleen, heart, skeletal muscle, pancreas, small intestine, and colon) from C57BL/6J male mice fed a low-fat diet (LFD) or a high-fat diet (HFD) for 12 weeks were also collected as we have previously described (129).

The *Ctrp10/C1ql2*-KO mice (C57BL/6NCrl-*C1ql2^em1(IMPC)Mbp^*/Mmucd; stock number 050587-UCD) were generated using the CRISPR-cas9 method at UC Davis. The two guide RNAs (gRNA) used were 5’-CCGGCGCC GCTCCACCATTACCT-3’ and 5’-TCAGGCCACCCCATCCCCATCGG-3’. The *Ctrp10* gene consists of 2 exons. The entire protein coding region spanning exon 1 and 2 was deleted. This KO strategy ensures a complete null allele for *Ctrp10*. The KO mice were maintained on a C57BL6/6J genetic background. Genotyping primers for WT allele were forward (m10-Com-F) 5’-TGTCGGGCTCTTCGACTCTCCA-3’ and reverse (m10-WT-R) 5’-GCATCTCGT AGTGAGCCGCTCC-3’. The size of the WT band was 360 bp. Genotyping primers for the *Ctrp10* KO allele were forward (m10-Com-F) 5’-TGTCGGGCTCTTCGACTCTCCA-3’ and reverse (m10-Mut-R1) 5’-GTCCAATCAGCT TTCTCAAGTCTGG-3’. The size of the KO band was 422 bp. The genotyping PCR parameters were as follows: 94°C for 5 min, followed by 10 cycles of (94°C for 10 sec, 65°C for 15 sec, 72°C for 30 sec), then 25 cycles of (94°C for 10 sec, 55°C for 15 sec, 72°C for 30 sec), and lastly 72°C for 5 min. Due to the presence of GC rich sequences, 7% DMSO was included in the PCR genotyping reaction. Mice were generated by intercrossing *Ctrp10* heterozygous (+/-) mice, supplemented with intercrossing WT or KO mice. *Ctrp10* KO (-/-) and WT (+/+) controls were housed in polycarbonate cages on a 12-h light–dark photocycle with ad libitum access to water and food. Mice were fed either a control low-fat diet (LFD; 10% kcal derived from fat; # D12450B; Research Diets, New Brunswick, NJ) or a high-fat diet (HFD; 60% kcal derived from fat; #D12492, Research Diets). LFD was provided for the duration of the study, beginning at 5 weeks of age; HFD was provided for 14 weeks, beginning at 6-7 weeks of age. At termination of study, all mice were fasted for 2 h and euthanized. Tissues were collected, snap-frozen in liquid nitrogen, and kept at −80°C until analysis. All mouse protocols (protocol # MO22M367) were approved by the Institutional Animal Care and Use Committee of the Johns Hopkins University School of Medicine. All animal experiments were conducted in accordance with the National Institute of Health guidelines and followed the standards established by the Animal Welfare Acts.

### Body composition analysis

Body composition analyses for total fat, lean mass, and water content were determined using a quantitative magnetic resonance instrument (Echo-MRI-100, Echo Medical Systems, Waco, TX) at the Mouse Phenotyping Core facility at Johns Hopkins University School of Medicine.

### Indirect calorimetry

LFD- or HFD-fed WT and *Ctrp10* KO male and female mice were used for simultaneous assessments of daily body weight change, food intake (corrected for spillage), physical activity, and whole-body metabolic profile in an open flow indirect calorimeter (Comprehensive Laboratory Animal Monitoring System, CLAMS; Columbus Instruments, Columbus, OH) as previously described (25). In brief, data were collected for three days to confirm mice were acclimatized to the calorimetry chambers (indicated by stable body weights, food intakes, and diurnal metabolic patterns), and data were analyzed from the fourth day. Rates of oxygen consumption (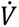_O2_; mL·kg^-1^·h^-1^) and carbon dioxide production (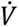_CO2_; mL·kg^-1·^h^-1^) in each chamber were measured every 24 min throughout the studies. Respiratory exchange ratio (RER 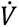_CO2_/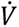_O2_) was calculated by CLAMS software (version 5.66) to estimate relative oxidation of carbohydrates (RER = 1.0) versus fats (RER = 0.7), not accounting for protein oxidation. Energy expenditure (EE) was calculated as EE= 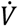_O2_× [3.815 + (1.232 × RER)] and normalized to lean mass. Because normalizing to lean mass can potentially lead to overestimation of EE, we also performed ANCOVA analysis on EE using body weight as a covariate (130). Physical activities were measured by infrared beam breaks in the metabolic chamber.

### Measurements of 24 h food intake

To independently confirm the food intake data collected in the metabolic cage (CLAMS), we also performed 24 h food intake measurements manually. All mice were singly housed, with wire mesh flooring inserts over a piece of cage paper on the bottom of the cage. A known weight of food pellets was given to each mouse. Twenty-four hours later, the leftover food pellets remaining on the flooring insert, along with any spilled crumbs on the cage paper, were collected and weighed. Thus, food intake was corrected for spillage.

### Glucose, insulin, pyruvate, and lipid tolerance tests

All tolerance tests were conducted as previously described (21, 24, 29). For glucose tolerance tests (GTTs), mice were fasted for 6 h before glucose injection. Glucose (Sigma, St. Louis, MO) was reconstituted in saline (0.9 g NaCl/L), sterile-filtered, and injected intraperitoneally (i.p.) at 1 mg/g body weight (i.e., 10 μL/g body weight). Blood glucose was measured at 0, 15, 30, 60, and 120 min after glucose injection using a glucometer (NovaMax Plus, Billerica, MA). For insulin tolerance tests (ITTs), food was removed 2 h before insulin injection. 6.5 μL of insulin stock (4 mg/mL; Gibco) was diluted in 10 mL of saline, sterile-filtered, and injected i.p. at 0.75 U/kg body weight (i.e., 10 μL/g body weight). Blood glucose was measured at 0, 15, 30, 60, and 90 min after insulin injection using a glucometer (NovaMax Plus).

### Fasting-Refeeding insulin tests

Mice were fasted overnight (∼16 h) then reintroduced to food as described (26). Blood glucose was monitored at the 16 h fast time point (time = 0 h refed) and at 1 and 2 hours into the refeeding process. Serum was collected at the 16 h fast and 2 h refed time points for insulin ELISA, as well as for the quantification of triglyceride, cholesterol, non-esterified free fatty acids (NEFA), and β-hydroxybutyrate levels.

### Blood and tissue chemistry analysis

Tail vein blood samples were allowed to clot on ice and then centrifuged for 10 min at 10,000 x *g*. Serum samples were stored at −80°C until analyzed. Serum triglycerides (TG) and cholesterol levels were measured according to manufacturer’s instructions using an Infinity kit (Thermo Fisher Scientific, Middletown, VA). Non-esterified free fatty acids (NEFA) were measured using a Wako kit (Wako Chemicals, Richmond, VA). Serum β-hydroxybutyrate (ketone) concentrations were measured with a StanBio Liquicolor kit (StanBio Laboratory, Boerne, TX). ELISA kits were used to measure serum insulin (Crystal Chem, Elk Grove Village, IL; cat # 90080), estradiol (Calbiotech, El Cajon, CA; cat # ES380S), and FGF21 (R&D Systems, Minneapolis, MN; cat # MF2100) according to the manufacturer’s instructions. Hydroxyproline assay (Sigma Aldrich, MAK008) was used to quantify total collagen content in liver and adipose tissues according to the manufacturer’s instructions. Lipid peroxidation levels (marker of oxidative stress) in the liver and adipose tissues were assessed by the quantification of malondialdehyde (MDA) via Thiobarbituric Acid Reactive Substances (TBARS) assay (Cayman Chemical, 700870) according to the manufacturer’s instructions.

### Serum lipoprotein triglyceride and cholesterol analysis by FPLC

Food was removed for 2-4 hr (in the light cycle) prior to blood collection. Sera collected from mice were pooled (*n* = 6-7/genotype) and sent to the Mouse Metabolism Core at Baylor College of Medicine for analysis. Serum samples were first fractionated by fast protein liquid chromatography (FPLC). A total of 45 fractions were collected, and TG and cholesterol in each fraction was quantified.

### Histology and quantification

Inguinal (subcutaneous) white adipose tissue (iWAT), gonadal (visceral) white adipose tissue (gWAT), and liver were dissected and fixed in formalin. Paraffin embedding, tissue sectioning, and staining with hematoxylin and eosin were performed at the Pathology Core facility at Johns Hopkins University School of Medicine. Images were captured with a Keyence BZ-X700 All-in-One fluorescence microscope (Keyence Corp., Itasca, IL). Adipocyte (gWAT and iWAT) cross-sectional area (CSA), as well as the total area covered by lipid droplets in hepatocytes were measured on hematoxylin and eosin-stained slides using ImageJ software (131). For CSA measurements, all cells in one field of view at 100X magnification per tissue section per mouse were analyzed. Image capturing and quantifications were carried out blinded to genotype.

### Fecal bomb calorimetry and assessment of fecal parameters

Fecal pellet frequency and average fecal pellet weight were monitored by housing each mouse singly in clean cages with a wire mesh sitting on top of a cutout cardboard that lay at the bottom of the cage for fecal collection. The number of fecal pellets and their total weight was recorded at the end of 24 h period. Additional fecal pellets collected for 3 full days were combined and shipped to the University of Michigan Animal Phenotyping Core for fecal bomb calorimetry. Briefly, fecal samples were dried overnight at 50°C prior to weighing and grinding them to powder. Each sample was mixed with wheat flour (90% wheat flour, 10% sample) and formed into 1.0 g pellet, which was then secured into the firing platform and surrounded by 100% oxygen. The bomb was lowered into a water reservoir and ignited to release heat into the surrounding water. These data were used to calculate fecal pellet frequency (bowel movements/day), average fecal pellet weight (g/bowel movement), fecal energy (cal/g feces), and total fecal energy (kcal/day).

### Mitochondrial respirometry of tissue samples

High-resolution respirometry was conducted on previously frozen tissue samples (liver and gastrocnemius muscle) to assay for mitochondrial activity as we have previously described (132). Briefly, all tissues were dissected, snap frozen in liquid nitrogen, and stored at −80°C for later analysis. Samples were thawed in MAS buffer (70mM sucrose, 220 mM mannitol, 5 mM KH_2_PO_4_, 5 mM MgCl_2_, 1 mM EGTA, 2 mM HEPES pH 7.4), finely minced with scissors, and then homogenized with a glass Dounce homogenizer. The resulting homogenate was spun at 1000 *g* for 10 min at 4°C. The supernatant was collected and immediately used for protein quantification by BCA assay (Thermo Scientific, 23225). Each well of the Seahorse microplate was loaded with 8 µg (liver samples) or 10 µg (gastrocnemius muscle samples) of homogenate protein. Each biological replicate is comprised of three technical replicates. Samples from all tissues were treated separately with NADH (1 mM) as a complex I substrate or Succinate (a complex II substrate, 5 mM) in the presence of rotenone (a complex I inhibitor, 2 µM), then with the inhibitors rotenone (2 µM) and Antimycin A (4 µM), followed by TMPD (0.45 mM) and Ascorbate (1 mM) to activate complex IV, and finally treated with Azide (40 mM) to assess non-mitochondrial respiration. All respiration data were normalized to mitochondrial content, quantified using MitoTracker Deep Red (MTDR, ThermoFisher, M22426) as described (132). Briefly, lysates were incubated with MTDR (1 µM) for 10 min at 37°C, then centrifuged at 2000 g for 5 min at 4°C. The supernatant was carefully removed and replaced with 1x MAS solution and fluorescence was read with excitation and emission wavelengths of 625 nm and 670 nm, respectively. To minimize non-specific background signal contribution, control wells were loaded with MTDR and 1x MAS and subtracted from all sample values.

### Tissue library preparation and RNA sequencing

Total RNA was isolated from tissues using Trizol reagent (Thermo Fisher Scientific) according to the manufacturer’s instructions. Library preparation and bulk RNA sequencing of liver, skeletal muscle (gastrocnemius), gonadal white adipose tissue (gWAT), and inguinal white adipose tissue (iWAT) of HFD-fed *Ctrp10*-KO female mice and WT controls were performed by Novogene (Sacramento, California, USA) on an Illumina platform (NovaSeq 6000) and pair-end reads were generated. Sample size: 6 WT and 6 KO for each tissue. All raw sequencing files are available from the NIH Sequence Read Archive (SRA) accession PRJNA971939.

### Mouse RNA-Sequencing analysis

Transcript features were assembled from raw fastq files and aligned to the current version of mouse transcriptome (Mus musculus.GRCm39.cdna) using kallisto-aln (133). Version-specific Ensembl transcript IDs were linked to gene symbols using biomart. Estimated counts were log normalized and filtered for a limit of sum >5 across all samples. Logistic regressions comparing WT vs KO samples across tissues were performed using limma (73). Differential expression results were visualized using available R packages in CRAN: ggplot2, ggVennDiagram and pheatmap. Gene set enrichment analyses of the DEGs were performed using Enrichr (134). Scripts for analyses and visualization are available at the github link below (Data availability section).

### Human sex difference analysis

All the datasets and scripts to perform analyses are available at the github link below (Data availability section). Male and Female human data were obtained from Genotype-Tissue Expression (GTEx) (86) and filtered for sufficient comparison of inter-tissue transcript correlation as described (87, 88). CTRP10/C1QL2 and other human orthologues for mouse differentially expressed genes (DEGs) were identified by intersecting mouse gene symbols with known human orthologues from the vertebrate homology resource at Mouse Genome Informatics (MGI) (135). Co-correlation between all human orthologues DEGs were calculated in either self-reported male or female subjects in GTEx using the bicorAndPvalue() function in Weighted Genetic Coexpression Network Analysis (WGCNA) package (136). To compare sex-differences of regression coefficients, Wilcoxon t-tests were compared between coefficients using the R package ggpubr. To investigate the degree of conservation of CTRP-engaged pathways, we mapped the DEGs identified from *Ctrp10* KO versus WT mice to their human orthologs, including human CTRP10/C1QL2, in the GTEx database for transcriptional correlations. Individuals were stratified by sex to examine sex-specific gene connectivity, consisting of 210 males and 100 females to compare gene expression across tissues. Gene-connectivity analyses were performed based on population correlation significances summarized by cumulative -log10(pvalues) as previously described (87, 137–139). Additional details on the filtering criteria and normalization methods can be found in the GitHub repository.

### Quantitative real-time PCR

Total RNA was isolated from tissues using Trizol reagent (Thermo Fisher Scientific). Purified RNA was reverse transcribed using an iScript cDNA Synthesis Kit (Bio-rad). Real-time quantitative PCR analysis was performed on a CFX Connect Real-Time System (Bio-rad) using iTaq^TM^ Universal SYBR Green Supermix (Bio-rad) according to manufacturer’s instructions. Data were normalized to the stable housekeeping gene *β-actin* or *36B4* (encoding the acidic ribosomal phosphoprotein P0) and expressed as relative mRNA levels using the ΔΔCt method (140). Real-time qPCR primers used to assess *Ctrp10* expression across mouse tissues were: *Ctrp10* forward, 5’-CGGCTTCATGAC ACTTCCTGA-3’ and reverse, 5’-AGCAGGGATGTGTCTTTTCCA-3’. qPCR primers used to confirm the absence of *Ctrp10* in KO mice were: forward (qPCR-m10-F2), 5’-CACGTACCACATTCTCATGCG-3’ and reverse (qPCR-m10-R1), 5’-TCGTAATTCTGGTCCGCGTC-3’.

### Statistical analyses

Sample size is indicated in figure and/or figure legend. All results are expressed as mean ± standard error of the mean (SEM). Statistical analysis was performed with Prism 9 software (GraphPad Software, San Diego, CA). Data were analyzed with two-tailed Student’s *t*-tests, one-way ANOVA or two-way ANOVA (with Sidak’s post hoc tests). 2-way ANOVA was used for body weight over time, fasting-refeeding response, and all tolerance tests. *P* < 0.05 was considered statistically significant.

## ACKNOWLEDGEMENTS

This work was supported by the National Institutes of Health (DK084171 to GWW, HL138193 and DK130640 to MMS). D.C.S. is supported by an NIH T32 training grant (HL007534). The fecal bomb calorimetry analysis was performed at the University of Michigan Animal Phenotyping Core, supported by center grants 1U2CDK135066-01 (Mi-MPMOD) and DK020572 (MDRC). The FPLC/serum analyses were performed the Mouse Metabolism and Phenotyping Core (MMPC) at the Baylor College of Medicine, supported by NIH grants (DK114356 and UM1HG006348).

## AUTHOR CONTRIBUTIONS

FC, GWW contributed to the experimental design; FC, DCS, MS, SA, and GWW performed the experiments; FC, DCS, MS, SA, LMV, MMS, and GWW analyzed and interpreted the data; GWW drafted the paper with inputs and edits from all authors.

## DATA AVAILABILITY

High-throughput sequencing data from this study have been submitted to the NCBI Sequence Read Archive under accession number PRJNA971939. All processed datasets used and R scripts to reproduce analyses are freely available at: https://github.com/Leandromvelez/CTRP10-Manuscript-DEG-Sex-specific-connectivities-and-integration/tree/main.

## ABBREVIATIONS

CTRP: C1q/TNF-related protein
DEG: Differentially expressed gene
EE: energy expenditure
gWAT: gonadal white adipose tissue
iWAT: inguinal white adipose tissue
HDL: High density lipoprotein
HFD: high-fat diet
i.p.: intraperitoneally
KO: knockout
LFD: low-fat diet
MHO: Metabolically healthy obese
NEFA: non-esterified free fatty acids
RER: respiratory exchange ratio
TG: triglyceride
VLDL: Very low density lipoprotein
WT: wildtype

## SUPPLEMENTAL FIGURE FILES LEGENDS AND SOURCE DATA

**Figure 4 – figure supplement 1.** No significant differences in serum estradiol levels between WT (*n* = 9) and *Ctrp10* KO (*n* = 6) female mice fed a low-fat diet (LFD). We included WT male sera (*n* = 5) as additional controls as chow-fed male mice are expected to have much lower levels of serum estradiol.

**Figure 4 – figure supplement 2.** Mitochondrial respiration through complex I (CI), CII, and CIV in liver and skeletal muscle (gastrocnemius) of WT (*n* = 9) and *Ctrp10* KO (*n* = 6) female mice fed a low-fat diet (LFD). No significant differences in liver and skeletal muscle (gastrocnemius) mitochondrial respiration through complex I (CI), CIII, and CIV between WT (*n* = 9) and *Ctrp10* KO (*n* = 6) female mice. (**A and D)** Average oxygen consumption rate (OCR) traces per group, normalized to mitochondrial content. Each group tracing represents the average trace of 9 WT and 6 KO samples. Each tracing shows the entire process of the Seahorse-based respirometry assay with injection compounds listed at the time of introduction to the sample. The sequence is as follows: i) basal reads, ii) addition of NADH (activation of respiration through complex I), iii) addition of antimycin A (AA, inhibitor of complex III) and rotenone (Rot, inhibitor of complex I), iv) addition of TMPD and ascorbate (to activate complex IV via electron donation to cytochrome c), and finally v) addition of azide (inhibitor of complex IV). **(B and E)** The same information as presented in (A and D) conducted on the same samples, but the NADH injection step is replaced with the injection of succinate (to activate respiration through complex II) and rotenone (to inhibit complex I). **(C and F)** Average values of all data presented for liver (A-B) and skeletal muscle (D-E). Each dot represents the average of three technical replicates measured at three separate times. Both independent measurements of complex IV (CIV) were used to determine average CIV respiration. * *p* < 0.05, *** *p* < 0.001, **** *p* < 0.0001 (two-way ANOVA with Fisher’s LSD test).

**Figure 5 – figure supplement 1.** No differences in food intake, body temperature, or changes in fecal output, frequency, and energy content between WT and *Ctrp10* KO female mice fed a high-fat diet. Sample size for food intake, fecal frequency, and fecal weight: WT = 6 and KO = 6; sample size for fecal energy and total fecal energy content: WT = 9 and KO = 6; sample size for deep colon temperature: WT = 9 and KO = 10.

**Figure 5 – figure supplement 2.** No significant differences in serum estradiol levels between WT (*n* = 6) and *Ctrp10* KO (*n* = 11) female mice fed a high-fat diet (HFD). We included WT male sera (*n* = 5) as additional controls as chow-fed male mice are expected to have much lower levels of serum estradiol.

**Figure 5 – figure supplement 3.** Mitochondrial respiration through complex I (CI), CII, and CIV in liver and skeletal muscle (gastrocnemius) of WT (*n* = 7) and *Ctrp10* KO (*n* = 8) female mice fed a high-fat diet. (**A and D)** Average oxygen consumption rate (OCR) traces per group, normalized to mitochondrial content. Each group tracing represents the average trace of 7 WT and 8 KO samples. Each tracing shows the entire process of the Seahorse-based respirometry assay with injection compounds listed at the time of introduction to the sample. The sequence is as follows: i) basal reads, ii) addition of NADH (activation of respiration through complex I), iii) addition of antimycin A (AA, inhibitor of complex III) and rotenone (Rot, inhibitor of complex I), iv) addition of TMPD and ascorbate (to activate complex IV via electron donation to cytochrome c), and finally v) addition of azide (inhibitor of complex IV). **(B and E)** The same information as presented in (A and D) conducted on the same samples, but the NADH injection step is replaced with the injection of succinate (to activate respiration through complex II) and rotenone (to inhibit complex I). **(C and F)** Average values of all data presented for liver (A-B) and skeletal muscle (D-E). Each dot represents the average of three technical replicates measured at three separate times. Both independent measurements of complex IV (CIV) were used to determine average CIV respiration. * *p* < 0.05, ** *p* < 0.01 (two-way ANOVA with Fisher’s LSD test).

**Figure 8 – source data 1.** Supplemental table with differentially expressed genes (DEGs) upregulated in the gonadal white adipose tissue (gWAT) of HFD-fed *Ctrp10* KO female mice relative to WT controls.

**Figure 8 – source data 2.** Supplemental table with differentially expressed genes (DEGs) down-regulated in the gonadal white adipose tissue (gWAT) of HFD-fed *Ctrp10* KO female mice relative to WT controls.

**Figure 8 – source data 3.** Supplemental table with differentially expressed genes (DEGs) upregulated in the inguinal white adipose tissue (iWAT) of HFD-fed *Ctrp10* KO female mice relative to WT controls.

**Figure 8 – source data 4.** Supplemental table with differentially expressed genes (DEGs) down-regulated in the inguinal white adipose tissue (iWAT) of HFD-fed *Ctrp10* KO female mice relative to WT controls.

**Figure 8 – source data 5.** Supplemental table with differentially expressed genes (DEGs) upregulated in the liver of HFD-fed *Ctrp10* KO female mice relative to WT controls.

**Figure 8 – source data 6.** Supplemental table with differentially expressed genes (DEGs) down-regulated in the liver of HFD-fed *Ctrp10* KO female mice relative to WT controls.

**Figure 8 – source data 7.** Supplemental table with differentially expressed genes (DEGs) upregulated in the skeletal muscle (gastrocnemius) of HFD-fed *Ctrp10* KO female mice relative to WT controls.

**Figure 8 – source data 8.** Supplemental table with differentially expressed genes (DEGs) down-regulated in the skeletal muscle (gastrocnemius) of HFD-fed *Ctrp10* KO female mice relative to WT controls.

**Figure 9 – figure supplement 1.** While hepatic *Fgf21* expression is significantly elevated, its systemic circulating levels were only found to be higher (*P* = 0.051) in LFD-fed, but no HFD-fed, KO female mice (Fig. 9 – figure supplement 1).

## REFERENCES

1. Collaboration, N. C. D. R. F. (2017) Worldwide trends in body-mass index, underweight, overweight, and obesity from 1975 to 2016: a pooled analysis of 2416 population-based measurement studies in 128.9 million children, adolescents, and adults. Lancet 390, 2627–2642

2. Flier, J. S. (2023) Moderating “the great debate”: The carbohydrate-insulin vs. the energy balance models of obesity. Cell Metab 35, 737–741

3. Bluher, M. (2019) Obesity: global epidemiology and pathogenesis. Nat Rev Endocrinol 15, 288–298

4. Bouchard, C. (2021) Genetics of Obesity: What We Have Learned Over Decades of Research. Obesity (Silver Spring) 29, 802–820

5. Loos, R. J. F., and Kilpelainen, T. O. (2018) Genes that make you fat, but keep you healthy. J Intern Med 284, 450–463

6. van Vliet-Ostaptchouk, J. V., Nuotio, M. L., Slagter, S. N., Doiron, D., Fischer, K., Foco, L., Gaye, A., Gogele, M., Heier, M., Hiekkalinna, T., Joensuu, A., Newby, C., Pang, C., Partinen, E., Reischl, E., Schwienbacher, C., Tammesoo, M. L., Swertz, M. A., Burton, P., Ferretti, V., Fortier, I., Giepmans, L., Harris, J. R., Hillege, H. L., Holmen, J., Jula, A., Kootstra-Ros, J. E., Kvaloy, K., Holmen, T. L., Mannisto, S., Metspalu, A., Midthjell, K., Murtagh, M. J., Peters, A., Pramstaller, P. P., Saaristo, T., Salomaa, V., Stolk, R. P., Uusitupa, M., van der Harst, P., van der Klauw, M. M., Waldenberger, M., Perola, M., and Wolffenbuttel, B. H. (2014) The prevalence of metabolic syndrome and metabolically healthy obesity in Europe: a collaborative analysis of ten large cohort studies. BMC Endocr Disord 14, 9

7. Bluher, M. (2020) Metabolically Healthy Obesity. Endocr Rev 41

8. Priest, C., and Tontonoz, P. (2019) Inter-organ cross-talk in metabolic syndrome. Nat Metab 1, 1177–1188

9. Seldin, M. M., Tan, S. Y., and Wong, G. W. (2014) Metabolic function of the CTRP family of hormones. Rev Endocr Metab Disord 15, 111–123

10. Wong, G. W., Wang, J., Hug, C., Tsao, T. S., and Lodish, H. F. (2004) A family of Acrp30/adiponectin structural and functional paralogs. Proc Natl Acad Sci U S A 101, 10302–10307

11. Seldin, M. M., Peterson, J. M., Byerly, M. S., Wei, Z., and Wong, G. W. (2012) Myonectin (CTRP15), a novel myokine that links skeletal muscle to systemic lipid homeostasis. J Biol Chem 287, 11968–11980

12. Wei, Z., Peterson, J. M., Lei, X., Cebotaru, L., Wolfgang, M. J., Baldeviano, G. C., and Wong, G. W. (2012) C1q/TNF-related protein-12 (CTRP12), a novel adipokine that improves insulin sensitivity and glycemic control in mouse models of obesity and diabetes. J Biol Chem 287, 10301–10315

13. Wei, Z., Peterson, J. M., and Wong, G. W. (2011) Metabolic regulation by C1q/TNF-related protein-13 (CTRP13): activation of AMP-activated protein kinase and suppression of fatty acid-induced JNK signaling. J Biol Chem 286, 15652–15665

14. Wei, Z., Seldin, M. M., Natarajan, N., Djemal, D. C., Peterson, J. M., and Wong, G. W. (2013) C1q/Tumor Necrosis Factor-related Protein 11 (CTRP11), a Novel Adipose Stroma-derived Regulator of Adipogenesis. J Biol Chem 288, 10214–10229

15. Wong, G. W., Krawczyk, S. A., Kitidis-Mitrokostas, C., Ge, G., Spooner, E., Hug, C., Gimeno, R., and Lodish, H. F. (2009) Identification and characterization of CTRP9, a novel secreted glycoprotein, from adipose tissue that reduces serum glucose in mice and forms heterotrimers with adiponectin. FASEB J 23, 241–258

16. Wong, G. W., Krawczyk, S. A., Kitidis-Mitrokostas, C., Revett, T., Gimeno, R., and Lodish, H. F. (2008) Molecular, biochemical and functional characterizations of C1q/TNF family members: adipose-tissue-selective expression patterns, regulation by PPAR-gamma agonist, cysteine-mediated oligomerizations, combinatorial associations and metabolic functions. Biochem J 416, 161–177

17. Ghai, R., Waters, P., Roumenina, L. T., Gadjeva, M., Kojouharova, M. S., Reid, K. B., Sim, R. B., and Kishore, U. (2007) C1q and its growing family. Immunobiology 212, 253–266

18. Ressl, S., Vu, B. K., Vivona, S., Martinelli, D. C., Sudhof, T. C., and Brunger, A. T. (2015) Structures of C1q-like proteins reveal unique features among the C1q/TNF superfamily. Structure 23, 688–699

19. Lei, X., Rodriguez, S., Petersen, P. S., Seldin, M. M., Bowman, C. E., Wolfgang, M. J., and Wong, G. W. (2016) Loss of CTRP5 improves insulin action and hepatic steatosis. Am J Physiol Endocrinol Metab 310, E1036–1052

20. Lei, X., Seldin, M. M., Little, H. C., Choy, N., Klonisch, T., and Wong, G. W. (2017) C1q/TNF-related protein 6 (CTRP6) links obesity to adipose tissue inflammation and insulin resistance. J Biol Chem 292, 14836–14850

21. Lei, X., and Wong, G. W. (2019) C1q/TNF-related protein 2 (CTRP2) deletion promotes adipose tissue lipolysis and hepatic triglyceride secretion. J Biol Chem 294, 15638–15649

22. Little, H. C., Rodriguez, S., Lei, X., Tan, S. Y., Stewart, A. N., Sahagun, A., Sarver, D. C., and Wong, G. W. (2019) Myonectin deletion promotes adipose fat storage and reduces liver steatosis. FASEB J 33, 8666–8687

23. Petersen, P. S., Lei, X., Wolf, R. M., Rodriguez, S., Tan, S. Y., Little, H. C., Schweitzer, M. A., Magnuson, T. H., Steele, K. E., and Wong, G. W. (2017) CTRP7 deletion attenuates obesity-linked glucose intolerance, adipose tissue inflammation, and hepatic stress. Am J Physiol Endocrinol Metab 312, E309–E325

24. Rodriguez, S., Lei, X., Petersen, P. S., Tan, S. Y., Little, H. C., and Wong, G. W. (2016) Loss of CTRP1 disrupts glucose and lipid homeostasis. Am J Physiol Endocrinol Metab 311, E678–E697

25. Sarver, D. C., Stewart, A. N., Rodriguez, S., Little, H. C., Aja, S., and Wong, G. W. (2020) Loss of CTRP4 alters adiposity and food intake behaviors in obese mice. Am J Physiol Endocrinol Metab 319, E1084–E1100

26. Sarver, D. C., Xu, C., Aja, S., and Wong, G. W. (2022) CTRP14 inactivation alters physical activity and food intake response to fasting and refeeding. Am J Physiol Endocrinol Metab 322, E480–E493

27. Sarver, D. C., Xu, C., Carreno, D., Arking, A., Terrillion, C. E., Aja, S., and Wong, G. W. (2022) CTRP11 contributes modestly to systemic metabolism and energy balance. FASEB J 36, e22347

28. Tan, S. Y., Lei, X., Little, H. C., Rodriguez, S., Sarver, D. C., Cao, X., and Wong, G. W. (2020) CTRP12 ablation differentially affects energy expenditure, body weight, and insulin sensitivity in male and female mice. Am J Physiol Endocrinol Metab 319, E146–E162

29. Tan, S. Y., Little, H. C., Lei, X., Li, S., Rodriguez, S., and Wong, G. W. (2016) Partial deficiency of CTRP12 alters hepatic lipid metabolism. Physiol Genomics 48, 936–949

30. Wei, Z., Lei, X., Petersen, P. S., Aja, S., and Wong, G. W. (2014) Targeted deletion of C1q/TNF-related protein 9 increases food intake, decreases insulin sensitivity, and promotes hepatic steatosis in mice. Am J Physiol Endocrinol Metab 306, E779–790

31. Wolf, R. M., Lei, X., Yang, Z. C., Nyandjo, M., Tan, S. Y., and Wong, G. W. (2016) CTRP3 deficiency reduces liver size and alters IL-6 and TGFbeta levels in obese mice. Am J Physiol Endocrinol Metab 310, E332–345

32. Peterson, J. M., Seldin, M. M., Tan, S. Y., and Wong, G. W. (2014) CTRP2 overexpression improves insulin and lipid tolerance in diet-induced obese mice. PLoS One 9, e88535

33. Peterson, J. M., Seldin, M. M., Wei, Z., Aja, S., and Wong, G. W. (2013) CTRP3 attenuates diet-induced hepatic steatosis by regulating triglyceride metabolism. Am J Physiol Gastrointest Liver Physiol 305, G214–224.

34. Peterson, J. M., Wei, Z., Seldin, M. M., Byerly, M. S., Aja, S., and Wong, G. W. (2013) CTRP9 transgenic mice are protected from diet-induced obesity and metabolic dysfunction. Am J Physiol Regul Integr Comp Physiol 305, R522–533

35. Appari, M., Breitbart, A., Brandes, F., Szaroszyk, M., Froese, N., Korf-Klingebiel, M., Mohammadi, M. M., Grund, A., Scharf, G. M., Wang, H., Zwadlo, C., Fraccarollo, D., Schrameck, U., Nemer, M., Wong, G. W., Katus, H. A., Wollert, K. C., Muller, O. J., Bauersachs, J., and Heineke, J. (2017) C1q-TNF-Related Protein-9 Promotes Cardiac Hypertrophy and Failure. Circ Res 120, 66–77

36. Zheng, Q., Yuan, Y., Yi, W., Lau, W. B., Wang, Y., Wang, X., Sun, Y., Lopez, B. L., Christopher, T. A., Peterson, J. M., Wong, G. W., Yu, S., Yi, D., and Ma, X. L. (2011) C1q/TNF-Related Proteins, A Family of Novel Adipokines, Induce Vascular Relaxation Through the Adiponectin Receptor-1/AMPK/eNOS/Nitric Oxide Signaling Pathway. Arterioscler Thromb Vasc Biol 31, 2616–2623

37. Kambara, T., Ohashi, K., Shibata, R., Ogura, Y., Maruyama, S., Enomoto, T., Uemura, Y., Shimizu, Y., Yuasa, D., Matsuo, K., Miyabe, M., Kataoka, Y., Murohara, T., and Ouchi, N. (2012) CTRP9 protein protects against myocardial injury following ischemia-reperfusion through AMP-activated protein kinase (AMPK)-dependent mechanism. J Biol Chem 287, 18965–18973

38. Kambara, T., Shibata, R., Ohashi, K., Matsuo, K., Hiramatsu-Ito, M., Enomoto, T., Yuasa, D., Ito, M., Hayakawa, S., Ogawa, H., Aprahamian, T., Walsh, K., Murohara, T., and Ouchi, N. (2015) C1q/Tumor Necrosis Factor-Related Protein 9 Protects against Acute Myocardial Injury through an Adiponectin Receptor I-AMPK-Dependent Mechanism. Mol Cell Biol 35, 2173–2185

39. Kanemura, N., Shibata, R., Ohashi, K., Ogawa, H., Hiramatsu-Ito, M., Enomoto, T., Yuasa, D., Ito, M., Hayakawa, S., Otaka, N., Murohara, T., and Ouchi, N. (2017) C1q/TNF-related protein 1 prevents neointimal formation after arterial injury. Atherosclerosis 257, 138–145

40. Ogawa, H., Ohashi, K., Ito, M., Shibata, R., Kanemura, N., Yuasa, D., Kambara, T., Matsuo, K., Hayakawa, S., Hiramatsu-Ito, M., Otaka, N., Kawanishi, H., Yamaguchi, S., Enomoto, T., Abe, T., Kaneko, M., Takefuji, M., Murohara, T., and Ouchi, N. (2020) Adipolin/CTRP12 protects against pathological vascular remodeling through suppression of smooth muscle cell growth and macrophage inflammatory response. Cardiovasc Res 116, 237–249

41. Otaka, N., Shibata, R., Ohashi, K., Uemura, Y., Kambara, T., Enomoto, T., Ogawa, H., Ito, M., Kawanishi, H., Maruyama, S., Joki, Y., Fujikawa, Y., Narita, S., Unno, K., Kawamoto, Y., Murate, T., Murohara, T., and Ouchi, N. (2018) Myonectin Is an Exercise-Induced Myokine That Protects the Heart From Ischemia-Reperfusion Injury. Circ Res 123, 1326–1338

42. Uemura, Y., Shibata, R., Ohashi, K., Enomoto, T., Kambara, T., Yamamoto, T., Ogura, Y., Yuasa, D., Joki, Y., Matsuo, K., Miyabe, M., Kataoka, Y., Murohara, T., and Ouchi, N. (2013) Adipose-derived factor CTRP9 attenuates vascular smooth muscle cell proliferation and neointimal formation. FASEB J 27, 25–33

43. Yuasa, D., Ohashi, K., Shibata, R., Mizutani, N., Kataoka, Y., Kambara, T., Uemura, Y., Matsuo, K., Kanemura, N., Hayakawa, S., Hiramatsu-Ito, M., Ito, M., Ogawa, H., Murate, T., Murohara, T., and Ouchi, N. (2016) C1q/TNF-related protein-1 functions to protect against acute ischemic injury in the heart. FASEB J 30, 1065–1075

44. Sun, Y., Yi, W., Yuan, Y., Lau, W. B., Yi, D., Wang, X., Wang, Y., Su, H., Gao, E., Koch, W. J., and Ma, X. L. (2013) C1q/Tumor Necrosis Factor-Related Protein-9, a Novel Adipocyte-Derived Cytokine, Attenuates Adverse Remodeling in the Ischemic Mouse Heart via Protein Kinase A Activation. Circulation 128, S113–120

45. Yi, W., Sun, Y., Yuan, Y., Lau, W. B., Zheng, Q., Wang, X., Wang, Y., Shang, X., Gao, E., Koch, W. J., and Ma, X. L. (2012) C1q/tumor necrosis factor-related protein-3, a newly identified adipokine, is a novel antiapoptotic, proangiogenic, and cardioprotective molecule in the ischemic mouse heart. Circulation 125, 3159–3169

46. Han, S., Jeong, A. L., Lee, S., Park, J. S., Buyanravjikh, S., Kang, W., Choi, S., Park, C., Han, J., Son, W. C., Yoo, K. H., Cheong, J. H., Oh, G. T., Lee, W. Y., Kim, J., Suh, S. H., Lee, S. H., Lim, J. S., Lee, M. S., and Yang, Y. (2018) C1q/TNF-alpha--Related Protein 1 (CTRP1) Maintains Blood Pressure Under Dehydration Conditions. Circ Res 123, e5–e19

47. Lee, S. M., Lee, J. W., Kim, I., Woo, D. C., Pack, C. G., Sung, Y. H., Baek, I. J., Jung, C. H., Kim, Y. H., and Ha, C. H. (2022) Angiogenic adipokine C1q-TNF-related protein 9 ameliorates myocardial infarction via histone deacetylase 7-mediated MEF2 activation. Sci Adv 8, eabq0898

48. Rodriguez, S., Little, H. C., Daneshpajouhnejad, P., Fenaroli, P., Tan, S. Y., Sarver, D. C., Delannoy, M., Talbot, C. C., Jr., Jandu, S., Berkowitz, D. E., Pluznick, J. L., Rosenberg, A. Z., and Wong, G. W. (2021) Aging and chronic high-fat feeding negatively affect kidney size, function, and gene expression in CTRP1-deficient mice. Am J Physiol Regul Integr Comp Physiol 320, R19–R35

49. Rodriguez, S., Little, H. C., Daneshpajouhnejad, P., Shepard, B. D., Tan, S. Y., Wolfe, A., Cheema, M. U., Jandu, S., Woodward, O. M., Talbot, C. C., Jr., Berkowitz, D. E., Rosenberg, A. Z., Pluznick, J. L., and Wong, G. W. (2020) Late-onset renal hypertrophy and dysfunction in mice lacking CTRP1. FASEB J 34, 2657–2676

50. Kirketerp-Moller, N., Bayarri-Olmos, R., Krogfelt, K. A., and Garred, P. (2020) C1q/TNF-Related Protein 6 Is a Pattern Recognition Molecule That Recruits Collectin-11 from the Complement System to Ligands. J Immunol 204, 1598–1606

51. Lahav, R., Haim, Y., Bhandarkar, N. S., Levin, L., Chalifa-Caspi, V., Sarver, D., Sahagun, A., Maixner, N., Kovesh, B., Wong, G. W., and Rudich, A. (2021) CTRP6 rapidly responds to acute nutritional changes, regulating adipose tissue expansion and inflammation in mice. Am J Physiol Endocrinol Metab 321, E702–E713

52. Ayyagari, R., Mandal, M. N., Karoukis, A. J., Chen, L., McLaren, N. C., Lichter, M., Wong, D. T., Hitchcock, P. F., Caruso, R. C., Moroi, S. E., Maumenee, I. H., and Sieving, P. A. (2005) Late-onset macular degeneration and long anterior lens zonules result from a CTRP5 gene mutation. Invest Ophthalmol Vis Sci 46, 3363–3371

53. Hayward, C., Shu, X., Cideciyan, A. V., Lennon, A., Barran, P., Zareparsi, S., Sawyer, L., Hendry, G., Dhillon, B., Milam, A. H., Luthert, P. J., Swaroop, A., Hastie, N. D., Jacobson, S. G., and Wright, A. F. (2003) Mutation in a short-chain collagen gene, CTRP5, results in extracellular deposit formation in late-onset retinal degeneration: a genetic model for age-related macular degeneration. Hum Mol Genet 12, 2657–2667

54. Luo, Y., Wu, X., Ma, Z., Tan, W., Wang, L., Na, D., Zhang, G., Yin, A., Huang, H., Xia, D., Zhang, Y., Shi, X., and Wang, L. (2016) Expression of the novel adipokine C1qTNF-related protein 4 (CTRP4) suppresses colitis and colitis-associated colorectal cancer in mice. Cell Mol Immunol 13, 688–699

55. Youngstrom, D. W., Zondervan, R. L., Doucet, N. R., Acevedo, P. K., Sexton, H. E., Gardner, E. A., Anderson, J. S., Kushwaha, P., Little, H. C., Rodriguez, S., Riddle, R. C., Kalajzic, I., Wong, G. W., and Hankenson, K. D. (2020) CTRP3 Regulates Endochondral Ossification and Bone Remodeling During Fracture Healing. J Orthop Res 38, 996–1006

56. Hamoud, N., Tran, V., Aimi, T., Kakegawa, W., Lahaie, S., Thibault, M. P., Pelletier, A., Wong, G. W., Kim, I. S., Kania, A., Yuzaki, M., Bouvier, M., and Cote, J. F. (2018) Spatiotemporal regulation of the GPCR activity of BAI3 by C1qL4 and Stabilin-2 controls myoblast fusion. Nature communications 9, 4470

57. Cho, Y., Kim, H. S., Kang, D., Kim, H., Lee, N., Yun, J., Kim, Y. J., Lee, K. M., Kim, J. H., Kim, H. R., Hwang, Y. I., Jo, C. H., and Kim, J. H. (2021) CTRP3 exacerbates tendinopathy by dysregulating tendon stem cell differentiation and altering extracellular matrix composition. Sci Adv 7, eabg6069

58. Kakegawa, W., Mitakidis, N., Miura, E., Abe, M., Matsuda, K., Takeo, Y. H., Kohda, K., Motohashi, J., Takahashi, A., Nagao, S., Muramatsu, S., Watanabe, M., Sakimura, K., Aricescu, A. R., and Yuzaki, M. (2015) Anterograde C1ql1 signaling is required in order to determine and maintain a single-winner climbing fiber in the mouse cerebellum. Neuron 85, 316–329

59. Sigoillot, S. M., Iyer, K., Binda, F., Gonzalez-Calvo, I., Talleur, M., Vodjdani, G., Isope, P., and Selimi, F. (2015) The Secreted Protein C1QL1 and Its Receptor BAI3 Control the Synaptic Connectivity of Excitatory Inputs Converging on Cerebellar Purkinje Cells. Cell reports 10, 820–832

60. Martinelli, D. C., Chew, K. S., Rohlmann, A., Lum, M. Y., Ressl, S., Hattar, S., Brunger, A. T., Missler, M., and Sudhof, T. C. (2016) Expression of C1ql3 in Discrete Neuronal Populations Controls Efferent Synapse Numbers and Diverse Behaviors. Neuron 91, 1034–1051

61. Sarver, D. C., Xu, C., Cheng, Y., Terrillion, C. E., and Wong, G. W. (2021) CTRP4 ablation impairs associative learning and memory. FASEB J 35, e21910

62. Matsuda, K., Budisantoso, T., Mitakidis, N., Sugaya, Y., Miura, E., Kakegawa, W., Yamasaki, M., Konno, K., Uchigashima, M., Abe, M., Watanabe, I., Kano, M., Watanabe, M., Sakimura, K., Aricescu, A. R., and Yuzaki, M. (2016) Transsynaptic Modulation of Kainate Receptor Functions by C1q-like Proteins. Neuron 90, 752–767

63. Huggett, S. B., and Stallings, M. C. (2020) Genetic Architecture and Molecular Neuropathology of Human Cocaine Addiction. J Neurosci 40, 5300–5313

64. Unroe, K. A., Glover, M. E., Shupe, E. A., Feng, N., and Clinton, S. M. (2021) Perinatal SSRI Exposure Disrupts G Protein-coupled Receptor BAI3 in Developing Dentate Gyrus and Adult Emotional Behavior: Relevance to Psychiatric Disorders. Neuroscience 471, 32–50

65. Sun, B. B., Chiou, J., Traylor, M., Benner, C., Hsu, Y. H., Richardson, T. G., Surendran, P., Mahajan, A., Robins, C., Vasquez-Grinnell, S. G., Hou, L., Kvikstad, E. M., Burren, O. S., Davitte, J., Ferber, K. L., Gillies, C. E., Hedman, A. K., Hu, S., Lin, T., Mikkilineni, R., Pendergrass, R. K., Pickering, C., Prins, B., Baird, D., Chen, C. Y., Ward, L. D., Deaton, A. M., Welsh, S., Willis, C. M., Lehner, N., Arnold, M., Worheide, M. A., Suhre, K., Kastenmuller, G., Sethi, A., Cule, M., Raj, A., Alnylam Human, G., AstraZeneca Genomics, I., Biogen Biobank, T., Bristol Myers, S., Genentech Human, G., GlaxoSmithKline Genomic, S., Pfizer Integrative, B., Population Analytics of Janssen Data, S., Regeneron Genetics, C., Burkitt-Gray, L., Melamud, E., Black, M. H., Fauman, E. B., Howson, J. M. M., Kang, H. M., McCarthy, M. I., Nioi, P., Petrovski, S., Scott, R. A., Smith, E. N., Szalma, S., Waterworth, D. M., Mitnaul, L. J., Szustakowski, J. D., Gibson, B. W., Miller, M. R., and Whelan, C. D. (2023) Plasma proteomic associations with genetics and health in the UK Biobank. Nature 622, 329–338

66. Kim, H., Bedsaul-Fryer, J. R., Schulze, K. J., Sincerbeaux, G., Baker, S., Rebholz, C. M., Wu, L. S., Gogain, J., Cuddeback, L., Yager, J. D., De Luca, L. M., Siddiqua, T. J., and West, K. P., Jr. (2024) An Early Gestation Plasma Inflammasome in Rural Bangladeshi Women. Biomolecules 14

67. Iijima, T., Miura, E., Watanabe, M., and Yuzaki, M. (2010) Distinct expression of C1q-like family mRNAs in mouse brain and biochemical characterization of their encoded proteins. Eur J Neurosci 31, 1606–1615

68. Vishvanath, L., and Gupta, R. K. (2019) Contribution of adipogenesis to healthy adipose tissue expansion in obesity. J Clin Invest 129, 4022–4031

69. Hotamisligil, G. S. (2006) Inflammation and metabolic disorders. Nature 444, 860–867

70. Sun, K., Tordjman, J., Clement, K., and Scherer, P. E. (2013) Fibrosis and adipose tissue dysfunction. Cell Metab 18, 470–477

71. Masschelin, P. M., Cox, A. R., Chernis, N., and Hartig, S. M. (2019) The Impact of Oxidative Stress on Adipose Tissue Energy Balance. Front Physiol 10, 1638

72. Hotamisligil, G. S. (2010) Endoplasmic reticulum stress and the inflammatory basis of metabolic disease. Cell 140, 900–917

73. Ritchie, M. E., Phipson, B., Wu, D., Hu, Y., Law, C. W., Shi, W., and Smyth, G. K. (2015) limma powers differential expression analyses for RNA-sequencing and microarray studies. Nucleic Acids Res 43, e47

74. Delezie, J., Dumont, S., Dardente, H., Oudart, H., Grechez-Cassiau, A., Klosen, P., Teboul, M., Delaunay, F., Pevet, P., and Challet, E. (2012) The nuclear receptor REV-ERBalpha is required for the daily balance of carbohydrate and lipid metabolism. FASEB J 26, 3321–3335

75. Hand, L. E., Usan, P., Cooper, G. J., Xu, L. Y., Ammori, B., Cunningham, P. S., Aghamohammadzadeh, R., Soran, H., Greenstein, A., Loudon, A. S., Bechtold, D. A., and Ray, D. W. (2015) Adiponectin induces A20 expression in adipose tissue to confer metabolic benefit. Diabetes 64, 128–136

76. Hunter, A. L., Pelekanou, C. E., Barron, N. J., Northeast, R. C., Grudzien, M., Adamson, A. D., Downton, P., Cornfield, T., Cunningham, P. S., Billaud, J. N., Hodson, L., Loudon, A. S., Unwin, R. D., Iqbal, M., Ray, D. W., and Bechtold, D. A. (2021) Adipocyte NR1D1 dictates adipose tissue expansion during obesity. Elife 10

77. Xu, J., Lloyd, D. J., Hale, C., Stanislaus, S., Chen, M., Sivits, G., Vonderfecht, S., Hecht, R., Li, Y. S., Lindberg, R. A., Chen, J. L., Jung, D. Y., Zhang, Z., Ko, H. J., Kim, J. K., and Veniant, M. M. (2009) Fibroblast growth factor 21 reverses hepatic steatosis, increases energy expenditure, and improves insulin sensitivity in diet-induced obese mice. Diabetes 58, 250–259

78. Sancar, G., Liu, S., Gasser, E., Alvarez, J. G., Moutos, C., Kim, K., van Zutphen, T., Wang, Y., Huddy, T. F., Ross, B., Dai, Y., Zepeda, D., Collins, B., Tilley, E., Kolar, M. J., Yu, R. T., Atkins, A. R., van Dijk, T. H., Saghatelian, A., Jonker, J. W., Downes, M., and Evans, R. M. (2022) FGF1 and insulin control lipolysis by convergent pathways. Cell Metab 34, 171–183 e176

79. Zhao, L., Fan, M., Zhao, L., Yun, H., Yang, Y., Wang, C., and Qin, D. (2020) Fibroblast growth factor 1 ameliorates adipose tissue inflammation and systemic insulin resistance via enhancing adipocyte mTORC2/Rictor signal. J Cell Mol Med 24, 12813–12825

80. Fan, L., Ding, L., Lan, J., Niu, J., He, Y., and Song, L. (2019) Fibroblast Growth Factor-1 Improves Insulin Resistance via Repression of JNK-Mediated Inflammation. Front Pharmacol 10, 1478

81. Kong, X., Feng, D., Wang, H., Hong, F., Bertola, A., Wang, F. S., and Gao, B. (2012) Interleukin-22 induces hepatic stellate cell senescence and restricts liver fibrosis in mice. Hepatology 56, 1150–1159

82. Yang, L., Zhang, Y., Wang, L., Fan, F., Zhu, L., Li, Z., Ruan, X., Huang, H., Wang, Z., Huang, Z., Huang, Y., Yan, X., and Chen, Y. (2010) Amelioration of high fat diet induced liver lipogenesis and hepatic steatosis by interleukin-22. J Hepatol 53, 339–347

83. Wang, X., Ota, N., Manzanillo, P., Kates, L., Zavala-Solorio, J., Eidenschenk, C., Zhang, J., Lesch, J., Lee, W. P., Ross, J., Diehl, L., van Bruggen, N., Kolumam, G., and Ouyang, W. (2014) Interleukin-22 alleviates metabolic disorders and restores mucosal immunity in diabetes. Nature 514, 237-241

84. Choi, C. S., Fillmore, J. J., Kim, J. K., Liu, Z. X., Kim, S., Collier, E. F., Kulkarni, A., Distefano, A., Hwang, Y. J., Kahn, M., Chen, Y., Yu, C., Moore, I. K., Reznick, R. M., Higashimori, T., and Shulman, G. I. (2007) Overexpression of uncoupling protein 3 in skeletal muscle protects against fat-induced insulin resistance. J Clin Invest 117, 1995–2003

85. Fan, L., Sweet, D. R., Prosdocimo, D. A., Vinayachandran, V., Chan, E. R., Zhang, R., Ilkayeva, O., Lu, Y., Keerthy, K. S., Booth, C. E., Newgard, C. B., and Jain, M. K. (2021) Muscle Kruppel-like factor 15 regulates lipid flux and systemic metabolic homeostasis. J Clin Invest 131

86. Consortium, G. T. (2013) The Genotype-Tissue Expression (GTEx) project. Nat Genet 45, 580–585

87. Velez, L. M., Van, C., Moore, T., Zhou, Z., Johnson, C., Hevener, A. L., and Seldin, M. M. (2022) Genetic variation of putative myokine signaling is dominated by biological sex and sex hormones. Elife 11

88. Sarver, D. C., Xu, C., Velez, L. M., Aja, S., Jaffe, A. E., Seldin, M. M., Reeves, R. H., and Wong, G. W. (2023) Dysregulated systemic metabolism in a Down syndrome mouse model. Mol Metab 68, 101666

89. Kloting, N., Fasshauer, M., Dietrich, A., Kovacs, P., Schon, M. R., Kern, M., Stumvoll, M., and Bluher, M. (2010) Insulin-sensitive obesity. Am J Physiol Endocrinol Metab 299, E506–515

90. Primeau, V., Coderre, L., Karelis, A. D., Brochu, M., Lavoie, M. E., Messier, V., Sladek, R., and Rabasa-Lhoret, R. (2011) Characterizing the profile of obese patients who are metabolically healthy. Int J Obes (Lond) 35, 971–981

91. Samocha-Bonet, D., Chisholm, D. J., Tonks, K., Campbell, L. V., and Greenfield, J. R. (2012) Insulin-sensitive obesity in humans - a ‘favorable fat’ phenotype? Trends Endocrinol Metab 23, 116–124

92. Stefan, N., Kantartzis, K., Machann, J., Schick, F., Thamer, C., Rittig, K., Balletshofer, B., Machicao, F., Fritsche, A., and Haring, H. U. (2008) Identification and characterization of metabolically benign obesity in humans. Arch Intern Med 168, 1609–1616

93. Calori, G., Lattuada, G., Piemonti, L., Garancini, M. P., Ragogna, F., Villa, M., Mannino, S., Crosignani, P., Bosi, E., Luzi, L., Ruotolo, G., and Perseghin, G. (2011) Prevalence, metabolic features, and prognosis of metabolically healthy obese Italian individuals: the Cremona Study. Diabetes Care 34, 210–215

94. Aguilar-Salinas, C. A., Garcia, E. G., Robles, L., Riano, D., Ruiz-Gomez, D. G., Garcia-Ulloa, A. C., Melgarejo, M. A., Zamora, M., Guillen-Pineda, L. E., Mehta, R., Canizales-Quinteros, S., Tusie Luna, M. T., and Gomez-Perez, F. J. (2008) High adiponectin concentrations are associated with the metabolically healthy obese phenotype. J Clin Endocrinol Metab 93, 4075–4079

95. Smith, G. I., Mittendorfer, B., and Klein, S. (2019) Metabolically healthy obesity: facts and fantasies. J Clin Invest 129, 3978–3989

96. Pataky, Z., Bobbioni-Harsch, E., and Golay, A. (2010) Open questions about metabolically normal obesity. Int J Obes (Lond) 34 **Suppl 2**, S18–23

97. Samuel, V. T., Petersen, K. F., and Shulman, G. I. (2010) Lipid-induced insulin resistance: unravelling the mechanism. Lancet 375, 2267–2277

98. Cohen, J. C., Horton, J. D., and Hobbs, H. H. (2011) Human fatty liver disease: old questions and new insights. Science 332, 1519–1523

99. Kim, J. Y., van de Wall, E., Laplante, M., Azzara, A., Trujillo, M. E., Hofmann, S. M., Schraw, T., Durand, J. L., Li, H., Li, G., Jelicks, L. A., Mehler, M. F., Hui, D. Y., Deshaies, Y., Shulman, G. I., Schwartz, G. J., and Scherer, P. E. (2007) Obesity-associated improvements in metabolic profile through expansion of adipose tissue. J Clin Invest 117, 2621–2637

100. Hotamisligil, G. S., Johnson, R. S., Distel, R. J., Ellis, R., Papaioannou, V. E., and Spiegelman, B. M. (1996) Uncoupling of obesity from insulin resistance through a targeted mutation in aP2, the adipocyte fatty acid binding protein. Science 274, 1377–1379

101. Brandon, A. E., Small, L., Nguyen, T. V., Suryana, E., Gong, H., Yassmin, C., Hancock, S. E., Pulpitel, T., Stonehouse, S., Prescott, L., Kebede, M. A., Yau, B., Quek, L. E., Kowalski, G. M., Bruce, C. R., Turner, N., and Cooney, G. J. (2022) Insulin sensitivity is preserved in mice made obese by feeding a high starch diet. Elife 11

102. Kusminski, C. M., Holland, W. L., Sun, K., Park, J., Spurgin, S. B., Lin, Y., Askew, G. R., Simcox, J. A., McClain, D. A., Li, C., and Scherer, P. E. (2012) MitoNEET-driven alterations in adipocyte mitochondrial activity reveal a crucial adaptive process that preserves insulin sensitivity in obesity. Nat Med 18, 1539–1549

103. Wang, F., Liu, H., Blanton, W. P., Belkina, A., Lebrasseur, N. K., and Denis, G. V. (2009) Brd2 disruption in mice causes severe obesity without Type 2 diabetes. Biochem J 425, 71–83

104. Karelis, A. D., Faraj, M., Bastard, J. P., St-Pierre, D. H., Brochu, M., Prud’homme, D., and Rabasa-Lhoret, R. (2005) The metabolically healthy but obese individual presents a favorable inflammation profile. J Clin Endocrinol Metab 90, 4145–4150

105. Sun, K., Kusminski, C. M., and Scherer, P. E. (2011) Adipose tissue remodeling and obesity. J Clin Invest 121, 2094–2101

106. Shulman, G. I. (2014) Ectopic fat in insulin resistance, dyslipidemia, and cardiometabolic disease. N Engl J Med 371, 1131–1141

107. Virtue, S., and Vidal-Puig, A. (2010) Adipose tissue expandability, lipotoxicity and the Metabolic Syndrome--an allostatic perspective. Biochim Biophys Acta 1801, 338–349

108. Samuel, V. T., and Shulman, G. I. (2012) Mechanisms for insulin resistance: common threads and missing links. Cell 148, 852–871

109. Summers, S. A., Chaurasia, B., and Holland, W. L. (2019) Metabolic Messengers: ceramides. Nat Metab 1, 1051–1058

110. Zhang, Y., Fang, B., Emmett, M. J., Damle, M., Sun, Z., Feng, D., Armour, S. M., Remsberg, J. R., Jager, J., Soccio, R. E., Steger, D. J., and Lazar, M. A. (2015) GENE REGULATION. Discrete functions of nuclear receptor Rev-erbalpha couple metabolism to the clock. Science 348, 1488–1492

111. Zhang, Y., Fang, B., Damle, M., Guan, D., Li, Z., Kim, Y. H., Gannon, M., and Lazar, M. A. (2016) HNF6 and Rev-erbalpha integrate hepatic lipid metabolism by overlapping and distinct transcriptional mechanisms. Genes Dev 30, 1636–1644

112. Yang, Y., Smith, D. L., Jr., Keating, K. D., Allison, D. B., and Nagy, T. R. (2014) Variations in body weight, food intake and body composition after long-term high-fat diet feeding in C57BL/6J mice. Obesity (Silver Spring) 22, 2147–2155

113. Rogers, N. H., Perfield, J. W., 2nd, Strissel, K. J., Obin, M. S., and Greenberg, A. S. (2009) Reduced energy expenditure and increased inflammation are early events in the development of ovariectomy-induced obesity. Endocrinology 150, 2161–2168

114. Hong, J., Stubbins, R. E., Smith, R. R., Harvey, A. E., and Nunez, N. P. (2009) Differential susceptibility to obesity between male, female and ovariectomized female mice. Nutr J 8, 11

115. Heine, P. A., Taylor, J. A., Iwamoto, G. A., Lubahn, D. B., and Cooke, P. S. (2000) Increased adipose tissue in male and female estrogen receptor-alpha knockout mice. Proc Natl Acad Sci U S A 97, 12729–12734

116. Bryzgalova, G., Lundholm, L., Portwood, N., Gustafsson, J. A., Khan, A., Efendic, S., and Dahlman-Wright, K. (2008) Mechanisms of antidiabetogenic and body weight-lowering effects of estrogen in high-fat diet-fed mice. Am J Physiol Endocrinol Metab 295, E904–912

117. Stubbins, R. E., Holcomb, V. B., Hong, J., and Nunez, N. P. (2012) Estrogen modulates abdominal adiposity and protects female mice from obesity and impaired glucose tolerance. Eur J Nutr 51, 861–870

118. Yonezawa, R., Wada, T., Matsumoto, N., Morita, M., Sawakawa, K., Ishii, Y., Sasahara, M., Tsuneki, H., Saito, S., and Sasaoka, T. (2012) Central versus peripheral impact of estradiol on the impaired glucose metabolism in ovariectomized mice on a high-fat diet. Am J Physiol Endocrinol Metab 303, E445–456

119. Mauvais-Jarvis, F., Clegg, D. J., and Hevener, A. L. (2013) The role of estrogens in control of energy balance and glucose homeostasis. Endocr Rev 34, 309–338

120. Correa, S. M., Newstrom, D. W., Warne, J. P., Flandin, P., Cheung, C. C., Lin-Moore, A. T., Pierce, A. A., Xu, A. W., Rubenstein, J. L., and Ingraham, H. A. (2015) An estrogen-responsive module in the ventromedial hypothalamus selectively drives sex-specific activity in females. Cell reports 10, 62–74

121. Butera, P. C. (2010) Estradiol and the control of food intake. Physiol Behav 99, 175–180

122. Xu, Y., Nedungadi, T. P., Zhu, L., Sobhani, N., Irani, B. G., Davis, K. E., Zhang, X., Zou, F., Gent, L. M., Hahner, L. D., Khan, S. A., Elias, C. F., Elmquist, J. K., and Clegg, D. J. (2011) Distinct hypothalamic neurons mediate estrogenic effects on energy homeostasis and reproduction. Cell Metab 14, 453–465

123. Krause, W. C., Rodriguez, R., Gegenhuber, B., Matharu, N., Rodriguez, A. N., Padilla-Roger, A. M., Toma, K., Herber, C. B., Correa, S. M., Duan, X., Ahituv, N., Tollkuhn, J., and Ingraham, H. A. (2021) Oestrogen engages brain MC4R signalling to drive physical activity in female mice. Nature 599, 131–135

124. Bolliger, M. F., Martinelli, D. C., and Sudhof, T. C. (2011) The cell-adhesion G protein-coupled receptor BAI3 is a high-affinity receptor for C1q-like proteins. Proc Natl Acad Sci U S A 108, 2534–2539

125. Sticco, M. J., Pena Palomino, P. A., Lukacsovich, D., Thompson, B. L., Foldy, C., Ressl, S., and Martinelli, D. C. (2021) C1QL3 promotes cell-cell adhesion by mediating complex formation between ADGRB3/BAI3 and neuronal pentraxins. FASEB J 35, e21194

126. Wang, J., Miao, Y., Wicklein, R., Sun, Z., Wang, J., Jude, K. M., Fernandes, R. A., Merrill, S. A., Wernig, M., Garcia, K. C., and Sudhof, T. C. (2021) RTN4/NoGo-receptor binding to BAI adhesion-GPCRs regulates neuronal development. Cell 184, 5869–5885 e5825

127. Alsharif, H., Latimer, M. N., Perez, K. C., Alexander, J., Rahman, M. M., Challa, A. K., Kim, J. A., Ramanadham, S., Young, M., and Bhatnagar, S. (2023) Loss of Brain Angiogenesis Inhibitor-3 (BAI3) G-Protein Coupled Receptor in Mice Regulates Adaptive Thermogenesis by Enhancing Energy Expenditure. Metabolites 13

128. Shiu, F. H., Wong, J. C., Bhattacharya, D., Kuranaga, Y., Parag, R. R., Alsharif, H. A., Bhatnagar, S., Van Meir, E. G., and Escayg, A. (2023) Generation and initial characterization of mice lacking full-length BAI3 (ADGRB3) expression. Basic Clin Pharmacol Toxicol

129. Rodriguez, S., Stewart, A. N., Lei, X., Cao, X., Little, H. C., Fong, V., Sarver, D. C., and Wong, G. W. (2019) PRADC1: a novel metabolic-responsive secretory protein that modulates physical activity and adiposity. FASEB J 33, 14748–14759

130. Tschop, M. H., Speakman, J. R., Arch, J. R., Auwerx, J., Bruning, J. C., Chan, L., Eckel, R. H., Farese, R. V., Jr., Galgani, J. E., Hambly, C., Herman, M. A., Horvath, T. L., Kahn, B. B., Kozma, S. C., Maratos-Flier, E., Muller, T. D., Munzberg, H., Pfluger, P. T., Plum, L., Reitman, M. L., Rahmouni, K., Shulman, G. I., Thomas, G., Kahn, C. R., and Ravussin, E. (2012) A guide to analysis of mouse energy metabolism. Nat Methods 9, 57–63

131. Schneider, C. A., Rasband, W. S., and Eliceiri, K. W. (2012) NIH Image to ImageJ: 25 years of image analysis. Nat Methods 9, 671–675

132. Sarver, D. C., Saqib, M., Chen, F., and Wong, G. W. (2024) Mitochondrial respiration atlas reveals differential changes in mitochondrial function across sex and age. Elife 13

133. Bray, N. L., Pimentel, H., Melsted, P., and Pachter, L. (2016) Near-optimal probabilistic RNA-seq quantification. Nat Biotechnol 34, 525–527

134. Kuleshov, M. V., Jones, M. R., Rouillard, A. D., Fernandez, N. F., Duan, Q., Wang, Z., Koplev, S., Jenkins, S. L., Jagodnik, K. M., Lachmann, A., McDermott, M. G., Monteiro, C. D., Gundersen, G. W., and Ma’ayan, A. (2016) Enrichr: a comprehensive gene set enrichment analysis web server 2016 update. Nucleic Acids Res 44, W90–97

135. Alliance of Genome Resources, C. (2020) Alliance of Genome Resources Portal: unified model organism research platform. Nucleic Acids Res 48, D650–D658

136. Langfelder, P., and Horvath, S. (2008) WGCNA: an R package for weighted correlation network analysis. BMC Bioinformatics 9, 559

137. Koplev, S., Seldin, M., Sukhavasi, K., Ermel, R., Pang, S., Zeng, L., Bankier, S., Di Narzo, A., Cheng, H., Meda, V., Ma, A., Talukdar, H., Cohain, A., Amadori, L., Argmann, C., Houten, S. M., Franzen, O., Mocci, G., Meelu, O. A., Ishikawa, K., Whatling, C., Jain, A., Jain, R. K., Gan, L. M., Giannarelli, C., Roussos, P., Hao, K., Schunkert, H., Michoel, T., Ruusalepp, A., Schadt, E. E., Kovacic, J. C., Lusis, A. J., and Bjorkegren, J. L. M. (2022) A mechanistic framework for cardiometabolic and coronary artery diseases. Nat Cardiovasc Res 1, 85–100

138. Massa, M. G., Scott, R. L., Cara, A. L., Cortes, L. R., Vander, P. B., Sandoval, N. P., Park, J. W., Ali, S. L., Velez, L. M., Wang, H. B., Ati, S. S., Tesfaye, B., Reue, K., van Veen, J. E., Seldin, M. M., and Correa, S. M. (2023) Feeding neurons integrate metabolic and reproductive states in mice. iScience 26, 107918

139. Zhou, M., Tamburini, I., Van, C., Molendijk, J., Nguyen, C. M., Chang, I. Y., Johnson, C., Velez, L. M., Cheon, Y., Yeo, R., Bae, H., Le, J., Larson, N., Pulido, R., Nascimento-Filho, C. H. V., Jang, C., Marazzi, I., Justice, J., Pannunzio, N., Hevener, A. L., Sparks, L., Kershaw, E. E., Nicholas, D., Parker, B. L., Masri, S., and Seldin, M. M. (2024) Leveraging inter-individual transcriptional correlation structure to infer discrete signaling mechanisms across metabolic tissues. Elife 12

140. Schmittgen, T. D., and Livak, K. J. (2008) Analyzing real-time PCR data by the comparative C(T) method. Nat Protoc 3, 1101–1108

141. Sievers, F., and Higgins, D. G. (2018) Clustal Omega for making accurate alignments of many protein sequences. Protein Sci 27, 135–145

